# Mitochondrial Transplantation: Adaptive Bio-enhancement

**DOI:** 10.1101/2024.09.20.614058

**Authors:** Xiaomeng Lu

## Abstract

Mitochondria, often referred to the powerhouse of the cell, are essential for cellular energy production, and their dysfunction can profoundly affect various organs. Transplantation of healthy mitochondria can restore the bioenergetics in diseased cells and address multiple conditions, but more potentials of this approach remain unclear. In this study, I demonstrated that the source of transplanted mitochondria is not limited by species, as exhibit no significant responses to mitochondria derived from different germlines. Moreover, I identified that metabolic compatibility between the recipient and exogenous mitochondria as a crucial factor in mitochondrial transplantation, which confers unique metabolic properties to recipient cells, enabling them to combat diseases. Additionally, my findings indicated competitive interactions among mitochondria with varying functions, with more bioenergetic-active mitochondria yielded superior therapeutic benefits. Notably, no upper limit for the bio-enhancement provided by exogenous mitochondria has been identified. Based on these insights, I proposes a novel therapeutic approach—adaptive bio-enhancement through mitochondrial transplantation.

## Introduction

Mitochondrial transplantation involves isolating mitochondria from normal cells or tissues and transferring them to diseased areas, either directly or indirectly, to enhance the function of damaged cells, thereby facilitating therapeutic interventions(1–3). Compared to traditional organ, tissue, or cell transplantation, mitochondrial transplantation offers a novel and less invasive alternative. In 2009, researchers at Boston Children ’s Hospital first reported this technique by injecting mitochondria into ischemic regions of the rabbit heart, resulting in significant therapeutic effects(4). Since then, mitochondrial transplantation has shown promising potential in treating various organ and tissue injuries, including those affecting the heart(5, 6), brain(7, 8), spinal cord(9), lungs, kidneys(10), liver(11), skeletal muscle(12), and skin(13). It has also been explored as a treatment for conditions such as sepsis(14) and cancer(15).

Despite the proven safety and therapeutic efficacy of mitochondrial transplantation in numerous diseases(16), its full potential remains to be fully elucidated. The transplanting of functionally normal mitochondria to restore the function of dysfunctional mitochondria in diseased cells involves differences in mitochondrial function, which can enhance the function of diseased cells(17–19). Furthermore, preemptively introducing exogenous mitochondria into healthy cells may increase their resistance to damage, thereby providing bioenergetic enhancement(6, 20). These insights raise a topic that warrants further investigation: the potential to select mitochondria with specific characteristics tailored to different disease populations or to employ mitochondria with superior functions instead of normal ones as an adaptive bio-enhancement strategy.

In this study, I expanded the sources of mitochondria for transplantation and demonstrated the broad applicability of this approach. By leveraging the germline-specific characteristics of mitochondria, I further treated cells under various disease conditions, highlighting the importance of metabolic matching. Additionally, hybrid mitochondria were generated with enhanced therapeutic effects by inducing cell fusion to combine multiple germline characteristics. Furthermore, I found that mitochondria with greater functional potency exerted markedly superior therapeutic effect compared to their germline-specific counterparts, even enhancing the function of normal cells to some extent. Similar results were observed in animal experiments, suggesting that mitochondrial transplantation could serve as a potential strategy for biological enhancement. These findings significantly expand the therapeutic potential of mitochondrial transplantation and open a new avenue for research into biological augmentation.

## Results

### Universality of Mitochondrial Transplantation

Current research on mitochondrial transplantation predominantly utilizes mitochondria from three sources: various cell lines(17–19, 21) such as human umbilical cord mesenchymal stem cells(22), and tissues such as liver(23), skeletal muscle(24), brain(25), and platelet-derived extracellular vesicles(26), The experimental models commonly employed include humans(27, 28), rabbits(4), mice(29), pigs(30), and rats(31). My study aimed to broaden the species diversity of mitochondria used for transplantation, thus enhance the therapeutic potential of this approach.

I obtained mitochondria from13 different types including african green monkey kidney cell(Vero), bovine kidney cell(MDCK), dog kidney cell(MDBK), cat kidney cell(CRFK), pig kidney cell(PK15), chicken liver cancer cell(LMH), spodoptera frugiperda cell(Sf9), turtle liver tissue, bullfrog liver tissue, sparrow liver tissue, lizard liver tissue, salmon liver tissue, and eel liver tissue, and assessed their functionally through adenosine triphosphate(ATP) assays (Figure 1A), JC-1 staining (Figure 1B), transmission electron microscopy (Figure 1C, Figure S1B), and mitochondrial respiratory chain complex activity assays (Figures 1D-G). My results confirmed that the isolated mitochondria were functionally intact and structurally well-preserved. Subsequently I co-cultured mitochondria from these species with human cardiomyocytes (AC16), human hepatoma cell (HepG2), and mouse fibroblast cell(L929), and efficient internalization as well as significant co-localization of the various mitochondria within recipient cells (Figure 1H, Figure S1A). To evaluate the safety of these transplantations, supernatants from the co-cultured cells were collected and their immune and inflammatory responses were assessed using enzyme-linked immunosorbent assay (ELISA). No significant changes were observed in the levels of IL-6, IL-10, and TNF-α in the mitochondria-transplanted groups compared to normal cells (Figures 1I-N), indicating a high degree of safety for multi-species mitochondrial transplantation.

**Figure 1:**
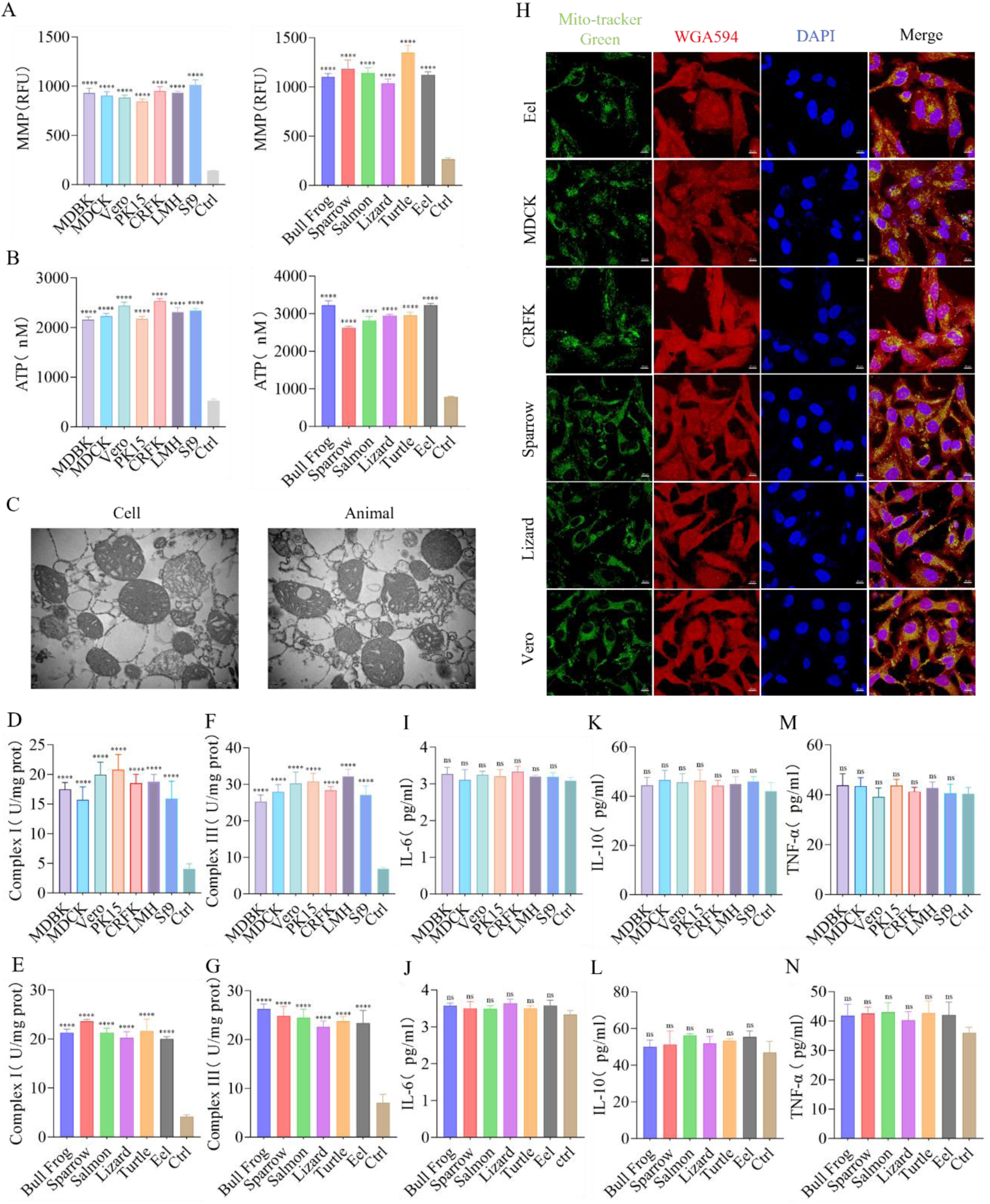
Universality of Mitochondrial Transplantation. (A) Analysis of membrane potential in both cells and animals isolated mitochondria. Ctrl: Mitochondria were subjected to two freeze-thaw cycles at -80°C and 37°C to completely disrupt their structure and function, serving as a comparison. n=3. (B) ATP production capacity in both cells and animals isolated mitochondria. Ctrl: Mitochondria were subjected to two freeze-thaw cycles at -80°C and 37°C, used for comparison. n=3. (C) Mitochondria were randomly selected from cells (Vero) and animals (Bull Frog) for electron microscopy to observe their morphology. Scale bar: 500 nm. (D-G) Activity of respiratory chain complexes I and III in both cells and animals isolated mitochondria. n=3. (H) Co-localization fluorescence imaging at 24 hours of transplanted mitochondria labeled with Mito-Tracker Green (green fluorescence) from Eel, MDCK, CRFK, Sparrow, Vero, and Lizard, alongside AC16, L929, and HepG2 cells labeled with WGA594 (red fluorescence). Blue fluorescence (DAPI) represents the nucleus. Scale bar: 10 μm. (I-N) Immunoreactivity of IL-6, IL-10, and TNF-α in cell culture medium collected 24 hours after the transplantation of exogenous mitochondria. n=3. Ctrl: Untreated group. ns = no statistical significance, **** = p<0.0001.

Additionally, a group of mitochondria from *Vaucheria litorea* was introduced to further investigate interspecies barriers. The mitochondria were labeled with Mito-Tracker Red and co-cultured with HepG2 cells pre-stained with WGA488 green fluorescence (Figures S2A-B). Due to limitations in the extraction technique, most *Vaucheria litorea* mitochondria were damaged. However, some of them with intact structure and function was successfully internalized. During the 25-day transplantation, no significant immune or inflammatory responses were recorded in the first 15 days. From day 16 onward, the contamination led to a rapid increase in IL-6, IL-10, and TNF-α levels, ultimately causing cell death (Figures S2D-H). These outcomes are acceptable because of the high contamination risk associated with plant mitochondrial isolation. The genomic differences between plant and animal mitochondria indicate substantial phylogenetic divergence(32, 33). Nevertheless, the early-stage safety observed following *Vaucheria litorea* mitochondrial transplantation strengthens my hypothesis, further supporting the universality of mitochondrial transplantation. Moreover, the internalization of mitochondria from multiple species demonstrates the cellular "inclusiveness" to various foreign organelles.

### Mitochondria with Matching Metabolic Characteristics Provide Improved Therapeutic Outcomes

Metabolic compatibility plays a crucial role in achieving optimal therapeutic outcomes during organ and stem cell transplantation, and this principle extends to mitochondrial transplantation(34–36). This raises a question: When mitochondrial functions are similar, do metabolically matched mitochondria offer better therapeutic outcomes? I hypothesized that metabolically matched mitochondria could yield superior therapeutic effects without significant functional differences. To minimize variables, mitochondria were selected from four different kidney cell species that were functionally similar, as determined by mitochondria membrane potential (MMP) and ATP assessments (Figures 2A, 2B, 2I, and 2J). These mitochondria were then co-cultured with cells under various disease model conditions, allowing us to assess varying therapeutic effects.

**Figure 2:**
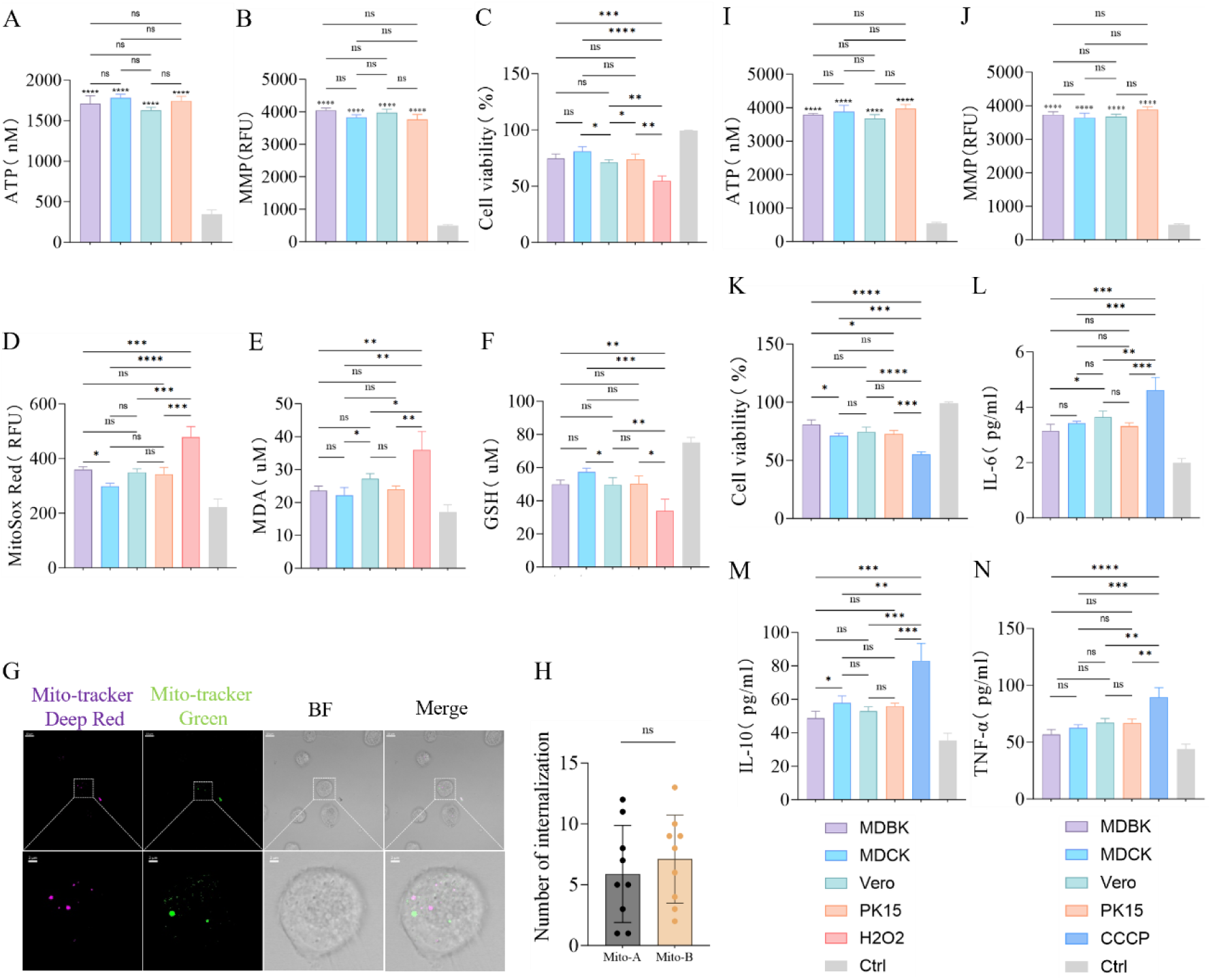
Mitochondria with Matching Metabolic Characteristics Provide Better Therapeutic Outcomes. (A, I) ATP production capacity in isolated mitochondria. Ctrl: Mitochondria subjected to two freeze-thaw cycles at -80° C and 37°C, used for comparison. n=3. (B, J) Membrane potential in isolated mitochondria. Ctrl: Mitochondria subjected to two freeze-thaw cycles at -80°C and 37°C, used for comparison. n=3. (C) Effects of mitochondrial transplantation from four different species on the viability of BMDM cells after H_2_O_2_ treatment. Ctrl: Untreated group. n=3. (D) Fluorescence imaging of mitochondrial ROS intensity in BMDM cells 24 hours after transplantation of mitochondria from four different species, followed by H_2_O_2_ treatment. Ctrl: Untreated group. n=3. (E-F) Effects of mitochondrial transplantation from four different species on MDA and GSH levels in BMDM cells after H_2_O_2_ treatment. Ctrl: Untreated group. n=3. (G-H) Fluorescence imaging and statistical analysis of the internalization of mitochondria A labeled with Mito-Tracker Red and mitochondria B labeled with Mito-Tracker Green in BMDM cells after H_2_O_2_ treatment. Original image scale bar: 10 µm; magnified region scale bar: 2 µm. n=9. (K) Effects of mitochondrial transplantation from four different species on the viability of AC16 cells after CCCP treatment. Ctrl: Untreated group. n=3. (L-N) Changes in IL-6, IL-10, and TNF-α levels in AC16 cells after CCCP treatment, following mitochondrial transplantation from four different species. Ctrl: Untreated group. n=3. ns = no statistical significance, * = p<0.05, ** = p<0.01, *** = p<0.001, **** = p<0.0001.

With the H_2_O_2_-induced Bone Marrow-Derived Macrophage (BMDM) oxidative stress model(37), all mitochondrial transplant groups showed significant therapeutic effects, but the MDCK group notably exhibited enhanced cell viability compared to the MDBK and Vero groups (Figure 2C). Fluorescence quantitative analysis revealed that mitochondrial ROS levels were significantly reduced in the MDCK group compared to those in the MDBK group (Figure 2D). Additionally, lipid peroxidation was significantly decreased, and glutathione levels were notably increased in the MDCK group compared to the Vero group (Figures 2E-F). Overall, these results suggest that MDCK mitochondrial transplantation is more effective in treating H_2_O_2_-induced oxidative stress in BMDMs. Then, I paired four types of mitochondria in pairs and labeled them with different fluorescent dyes, and co-culturing them with H_2_O_2_-treated BMDMs for 24 hours, fluorescence immunoimaging revealed no significant competition between the different mitochondria, and their internalization amounts in the recipient cells were similar (Figures 2G-H). According to these results, I inferred that the key distinguishing factor was their species of origin. Although the primary function of mitochondria is to provide energy, their auxiliary functions, which vary among species, appear to have made the MDCK mitochondria more metabolically compatible with the H_2_O_2_-treated BMDMs, resulting in improved therapeutic effects.

The results of mitochondrial transplantation from CCCP-treated AC16 cells further supported the hypothesis(38). Despite comparable mitochondrial function assessed by ATP and MMP assessments (Figures 2I-J), the MDBK group did not demonstrate a significant advantage. Nevertheless, its therapeutic effect was superior, markedly enhancing cell viability compared to the MDCK and PK15 groups (Figure 2K). Additionally, these four mitochondrial transplantation groups significantly reduced the inflammatory response, the MDBK group exhibited lower IL-6 levels compared to the Vero group (Figure 2L) and reduced IL-10 levels compared to the MDCK group (Figure 2M), although no significant differences were observed in TNF-α levels among four transplanted groups (Figure 2N). Overall, mitochondria derived from MDBK cells displayed enhanced anti-inflammatory effects in CCCP-treated AC16 cells. Despite lacking functional advantages, the improved therapeutic outcomes observed in the MDBK group suggest that its mitochondria were more metabolically compatible with the CCCP-treated AC16 cells. Similar results were also verified in the subsequent HepG2 cell model processed with LPS (Figure S3A-F).

### Mitochondrial Fusion Generates More Potent Pluripotent Hybrid Mitochondria

Mitochondrial transplantation can endow recipient cells with the characteristics of donor mitochondria. For instance, co-culturing mitochondria derived from human mammary epithelial cells with breast cancer cells has been demonstrated to inhibit glycolysis, reduce glucose uptake, suppress cancer cell proliferation, and enhance sensitivity to paclitaxel(39). Similarly, co-culturing mitochondria that contain antibiotic resistance genes with antibiotic-sensitive cells can bolster resistance to antibiotics(40). Given that cell fusion can lead to metabolic reprogramming, enabling fused cells to acquire traits from different cell types(41), a question emerges: Can the mitochondria within fused cells obtain multi-germline characteristics? Furthermore, could mitochondria possessing multi-germline traits offer enhanced therapeutic efficacy under equivalent functional conditions?

Specifically, HL1 and H9C2 cells were fused, and extracted for analysis using immunofluorescence and flow cytometry (Figures 3A-B). Then, the resulting cells were cultured and analyzed by Western blot on the second day, revealing a significantly higher expression of Opa1 in the HL1+H9C2 hybrid mitochondria compared to either HL1 or H9C2. Additionally, Drp1 and Mfn1 expression levels were notably elevated compared to those in HL1, with no significant difference observed compared to H9C2 (Figures 3C-D). Overall, the functional evaluation suggested that HL1+H9C2 hybrid mitochondria exhibited enhanced dynamic activity, indicating an active state of mitochondrial fusion within the cells. The three types of cells were further cultured until the fourth generation to ensure functional stability. Subsequently, oxidative stress was induced in AC16 cells with H₂O₂, and mitochondria were extracted from HL1, H9C2, and HL1+H9C2 cells for mitochondrial function evaluation through measurement of MMP and ATP levels. The results demonstrated no significant functional differences among these three types of mitochondria, all of which maintained normal functionality (Figures 3E-F). In addition, these mitochondria were co-cultured with AC16 cells. The AC16 cell viability was significantly enhanced within all groups, with the HL1+H9C2 group showing the most pronounced effect (Figure 3G). Quantitative fluorescence analysis revealed a significant reduction in mitochondrial ROS levels across all groups, with the HL1+H9C2 group indicating superior oxidative stress mitigation compared to the H9C2 group (Figure 3H). Notably, no significant differences in malondialdehyde (MDA) levels were observed among the three groups (Figure 3I). However, glutathione (GSH) levels in the HL1+H9C2 group were significantly higher than those in the H9C2 group (Figure 3J). These findings indicate that hybrid mitochondria derived from HL1+H9C2 could provide superior therapeutic effects on AC16 cells compared to the other two mitochondrial types.

**Figure 3:**
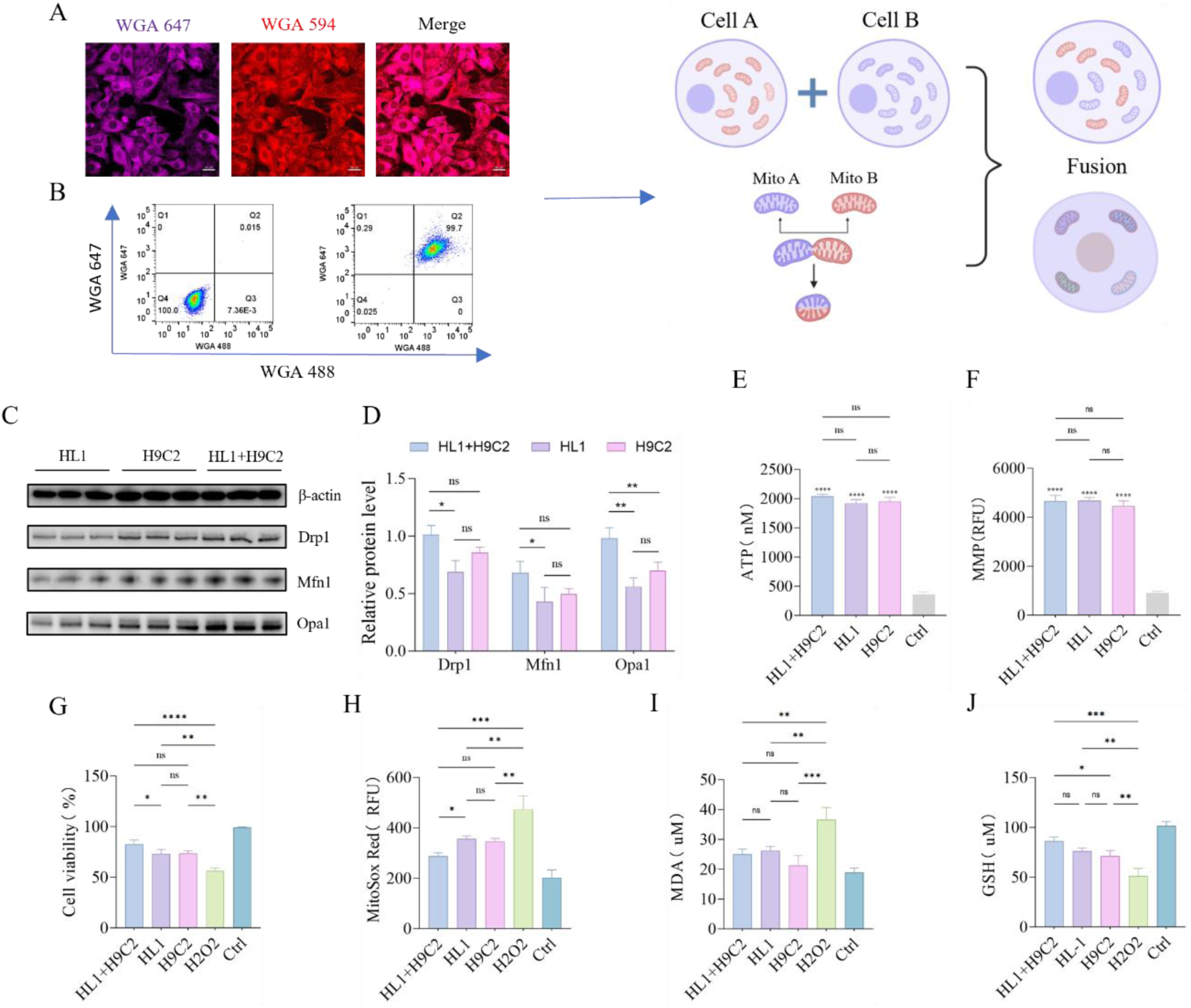
Generation of More Potent Pluripotent Hybrid Mitochondria through Mitochondrial Fusion. (A) Co-localization fluorescence imaging 24 hours after the fusion of HL1 cells labeled with WGA647 and H9C2 cells labeled with WGA594. Scale bar: 20 µm. (B) Flow cytometric sorting of hybrid cells 24 hours after the fusion of HL1 cells labeled with WGA647 and H9C2 cells labeled with WGA488. (C-D) Western blot analysis of the mitochondrial fusion and fission proteins Mfn1, Opa1, and Drp1 in HL1, H9C2, and HL1+H9C2 cells on the second day after cell fusion. n=3. (E) ATP production capacity in isolated mitochondria, Ctrl: Mitochondria subjected to two freeze-thaw cycles at -80°C and 37°C, used for comparison. n=3. (F) JC-1 membrane potential detection of isolated mitochondria. Ctrl: Mitochondria subjected to two freeze-thaw cycles at -80°C and 37°C, used for comparison. n=3. (G) Effects of transplantation of three different types of mitochondria on the viability of AC16 cells after H_2_O_2_ treatment. Ctrl: Untreated group. n=3. (H) Effects of transplantation of three different types of mitochondria on mitochondrial ROS levels in AC16 cells after H_2_O_2_ treatment. Ctrl: Untreated group. n=3. (I-J) Effects of transplantation of mitochondria from three different species on MDA and GSH levels in AC16 cells after H_2_O_2_ treatment. Ctrl: Untreated group. n=3. ns = no statistical significance, * = p<0.05, ** = p<0.01, *** = p<0.001, **** = p<0.0001

### Competitive Internalization of Mitochondria: The Fittest Functions Best

Natural selection operates on the principle that the fittest organisms are more likely to survive (42), a principle that also applies to mitochondrial transplantation. I think that the significantly differences in the bioenergetic function of transplanted mitochondria may allow more suitable mitochondria to provide better therapeutic outcomes.

Given the variability in bioenergetic function attributable to differences in the physiological state and the rearing environment of animals, I extracted mitochondria from the liver tissues of sparrows, lizards, salmon, and bullfrogs and co-cultured them with CCCP-treated HepG2 cells(43). Subsequently ATP and MMP assessments indicated that the mitochondria from bullfrogs exhibited the strongest bioenergetic function among the four groups (Figures 4A-B). The cell viability of HepG2 was improved within all groups, with the Bull Frog group showing a more pronounced effect compared to the Sparrow and Lizard groups (Figure 4C). Furthermore, the mitochondria ROS production was markedly reduced within the Bull Frog group compared to both Sparrow and Salmon groups (Figure 4D). The Bull Frog group also exhibited greater efficacy in repairing DNA oxidative damage than Sparrow and Salmon groups (Figure 4E). Immunofluorescence further revealed a significant competition between mitochondria with different functions, HepG2 cells internalized a higher quantity of bullfrog derived mitochondria than those derived from lizards (Figures 4F-G). These findings indicate that although metabolic compatibility is essential for similar bioenergetic function, significant improved bioenergetic performance offers greater therapeutic benefits in case of substantial disparity. That is to say, mitochondria with stronger overall function are more suitable for treatment.

**Figure 4:**
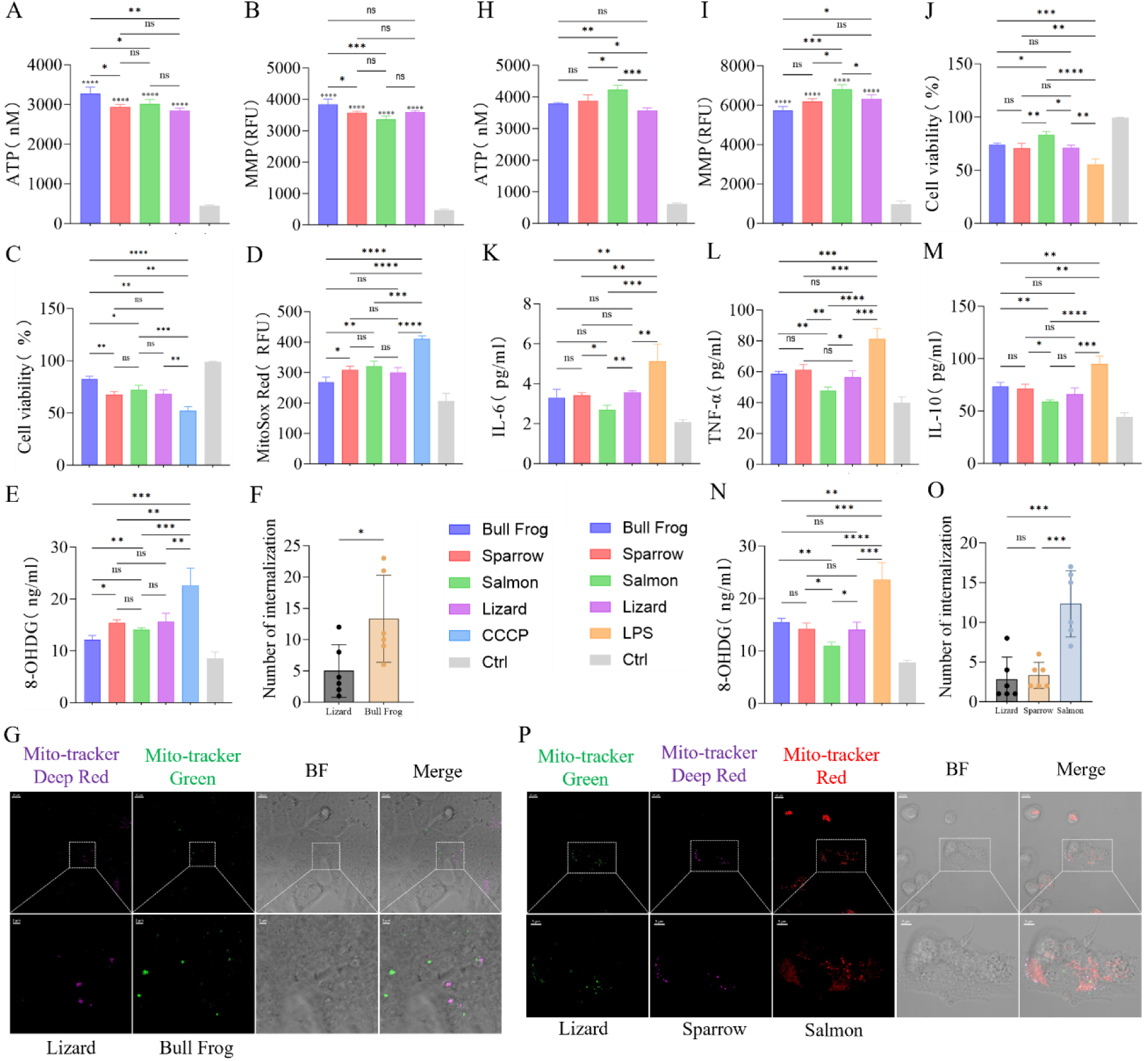
Competitive Internalization of Mitochondria: The Fittest Functions Best. (A, H) ATP production capacity in isolated mitochondria. Ctrl: Mitochondria subjected to two freeze-thaw cycles at - 80°C and 37°C, used for comparison. n=3. (B, I) Membrane potential in isolated mitochondria. Ctrl: Mitochondria subjected to two freeze-thaw cycles at -80°C and 37°C, used for comparison. n=3. (C) Effects of mitochondria transplantation from four different species on the viability of HepG2 cells after CCCP treatment. Ctrl: Untreated group. n=3. (D) Effects of mitochondria transplantation from four different species on mitochondria ROS in HepG2 cells after CCCP treatment. Ctrl: Untreated group. n=3. (E) Detection of DNA damage repair in HepG2 cells after CCCP treatment following mitochondria transplantation from four different species. Ctrl: Untreated group. n=3. (F-G) Fluorescence imaging and statistical analysis of the internalization of Lizard mitochondria labeled with Mito-Tracker Deep-Red and Bull Frog mitochondria labeled with Mito-Tracker Green in HepG2 cells after CCCP treatment. Original image scale bar: 10µm, magnified region scale bar: 2µm. n=6. (J) Effects of mitochondria transplantation from four different species on the viability of BMDM cells after LPS treatment. Ctrl: Untreated group. n=3. (K-M) Changes in IL-6, IL-10, and TNF-α levels in BMDM cells after LPS treatment following mitochondria transplantation from four different species. Ctrl: Untreated group. n=3. (N) Detection of DNA damage repair in BMDM cells after LPS treatment following mitochondria transplantation from four different species. n=3. (O-P) Fluorescence imaging and statistical analysis of the internalization of Lizard mitochondria labeled with Mito-Tracker Green, Sparrow mitochondria labeled with Mito-Tracker Deep-Red, and Salmon mitochondria labeled with Mito-Tracker Red in BMDM cells after LPS treatment. Original image scale bar: 10µm, magnified region scale bar: 5µm. n=6. ns = no statistical significance, * = p<0.05, ** = p<0.01, *** = p<0.001, **** = p<0.0001

Next, I extracted mitochondria from the liver tissues of the four animals and co-culturing them with LPS-treated BMDM cells(44). Assessments of ATP and MMP demonstrated that salmon derived mitochondria exhibited superior bioenergetic function compared to the other three groups (Figures 4H-I). Leveraging their robust biological effects, salmon mitochondria displayed enhanced therapeutic outcomes. Specifically, there were effective in increasing the viability of BMDM, reducing inflammation, and mitigating DNA oxidative damage. The Salmon group showed higher cell viability than the other three groups (Figure 4J), lower IL-6 levels compared to the Sparrow and Lizard groups (Figure 4K), lower TNF-α levels than the other three groups (Figure 4L), lower IL-10 levels than the Sparrow and Bull Frog groups (Figure 4M), and lower 8-OHDG levels than the other three groups (Figure 4N). Immunofluorescence analysis revealed that the Salmon group exhibited a dominant advantage, with its internalization quantity far exceeding that of the Lizard and Sparrow groups (Figures 4O-P). Similar results were observed in the D-gal-treated HepG2 cells model (Figures S3G-L). These findings indicate that, under equal conditions, the mitochondria compete to out- survive each other, and cells will preferentially select mitochondria that are more beneficial to them, demonstrating the classic law of "survival of the fittest."

### Mitochondrial Transplantation Enhances Physical Function and Biological Potency in Mice

The in vitro experiments indicated that mitochondria with greater functional potency exhibited superior therapeutic effects on diseases compared to mitochondria with normal function. I want to know in vivo experiments are equally applicable. Previous studies have suggested that preemptive administration of normally functioning mitochondria can enhance cellular resistance to stimuli(45), but the extent of its benefits to general recipients remains unclear. Mitochondrial function improved by training mitochondrial biogenesis (46, 47). Therefore, mice with trained mitochondria may exhibit better therapeutic outcomes than those with autologous or allogeneic mitochondria with normal function. The functional enhancement of mitochondria in trained mice is based on their overall physical fitness. Natural wild mice, being more robust with stronger physical fitness and metabolism, are hypothesized to possess even more potent mitochondrial function.

By isolating mitochondria from young C57BL/6J and wild mice, I conducted tail vein injections in mice with an acute inflammation model and normal ones. Additionally, I co-cultured these mitochondria with normal and LPS-treated BMDMs. ATP and MMP analyses revealed that the mitochondria derived from wild mice exhibited the highest functional potency (Figures 5A-B), with no immune inflammatory response observed after a 24 hours period (Figure S4B). Small animal imaging detected the absorption of exogenous mitochondria by severely damaged liver and kidney tissues through the circulatory system (Figure 5C). At three days post-transplantation, grip strength and endurance were significantly improved in all acute inflammation model groups, with the Wild Mice group showing the most notable endurance improvement (Figures 5D and 5I). In addition, no significant differences in grip strength were observed among the normal model groups, while the Wild Mice group demonstrated a substantial increase in endurance compared to the Ctrl group (Figures 5E and 5I). After one week, all acute inflammation model groups exhibited recovery in body weight, with the Wild Mice group demonstrating a significant increase compared to the LPS-Ctrl group (Figures 5G-H). In contrast, no significant differences in body weight were observed in the normal model groups (Figures 5F and 5H). MRI assessment revealed substantial increases in hind limb muscle volume for both the Wild Mice and Young C57 groups within the acute inflammation model, with the Wild Mice group showing slightly outperforming the Young C57 group (Figures S4D and 4E). Additionally, the muscle tissue damage recovery was also more effective in the Wild Mice group (Figures S4D and 4E). After treatment, serum levels of IL-6, IL-10, and TNF-α were reduced in both the Wild Mice and Young C57 groups of acute inflammation model, with the Wild Mice group showing lower levels of IL-10 and TNF-α compared to the Young C57 group, and no significant changes were observed in the normal model groups (Figures 6A-C). Furthermore, liver and kidney function were significantly improved in the Wild Mice and Young C57 groups of acute inflammation model, with serum AST levels significantly lower in the Wild Mice group compared to the Young C57 group. No significant differences were observed in the normal model groups (Figures 6D-F). Both the Wild Mice and Young C57 groups significantly reduced oxidative stress within the acute inflammation model. Characterized by decreased levels of MDA and increased superoxide dismutase (SOD) activity in the Wild Mice group compared to the Young C57 group. Moreover, the Wild Mice group demonstrated a notable increase in SOD activity among normal model groups. However, no difference in MDA levels were observed across the normal model groups (Figures 6G-H). Mitochondrial transplantation significantly reduced the expression of inflammatory proteins in the liver and kidney in the acute inflammation model group, with the Wild Mice group showing lower P-JNK protein expression compared to the Young C57 group (Figures 6I-K). Real-time quantitative PCR revealed a downregulation of IL-6, IL-8, IL-18, TGF-β, MPO, CXCL1, CXCL2, and HMGB1 as well as upregulation of IL-13, SOD1, ARG1, and COX1 in the kidneys of both Wild Mice and Young C57 groups when compared to the LPS-Ctrl group. In the liver, IL-6, IL-8, IL-18, IL-1β, CXCL2, CCL5, CCL10, and TGF-β were downregulated, while IL-4, IL-13, ARG1, and SOD1 were upregulated (Figures S4F-G). Notably, significant differences were identified between the Wild Mice and Young C57 groups, indicating that wild mice mitochondria could exert a superior therapeutic effect.

**Figure 5:**
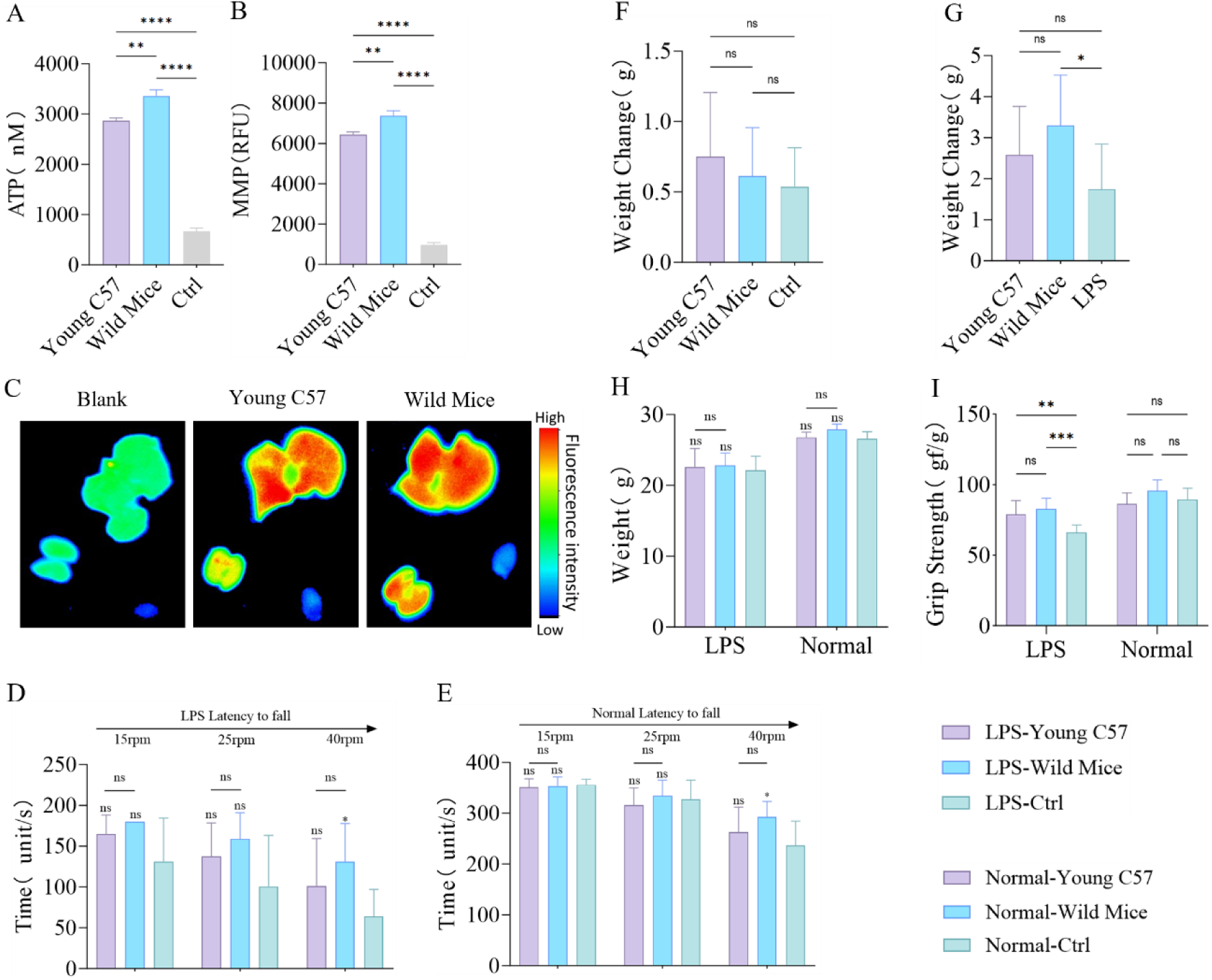
Mitochondrial Transplantation Enhances Physical Function and Biological Potency in Mice. (A-B) Evaluation of isolated mitochondrial functions through ATP production and mitochondrial membrane potential (MMP) assays, with mitochondria subjected to two freeze-thaw cycles at -80°C and 37°C to completely disrupt their structure and function as a control. n=3. (C) Biodistribution of exogenous mitochondria in the mouse liver, kidney, and heart detected by in vivo imaging 24 hours after intravenous tail vein injection of mitochondria. The blank group consists of untreated LPS-Ctrl group mice. (D-E) Assessment of motor function in acute inflammation model mice and normal model mice three days after the transplantation of two different types of mitochondria, with untreated groups as control. n=6. (F-H) Changes in body weight of acute inflammation model mice and normal model mice before and one week after the transplantation of two different types of mitochondria, with untreated groups as control. n=6. (I) Grip strength measurements in acute inflammation model mice and normal model mice three days after the transplantation of two different types of mitochondria, with untreated groups as control. n=6. ns = no statistical significance, * = p<0.05, ** = p<0.01, *** = p<0.001, **** = p<0.0001

**Figure 6:**
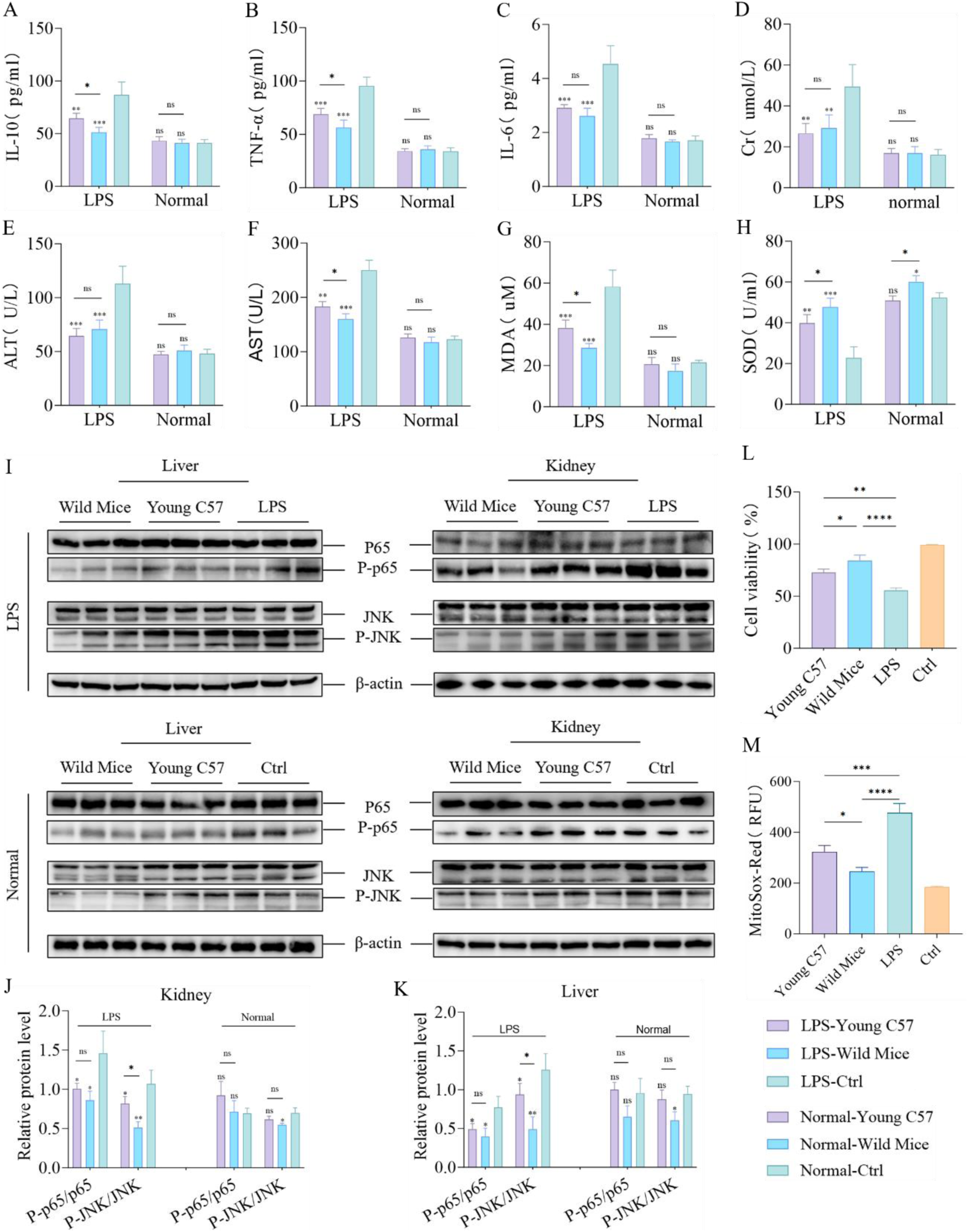
Mitochondrial Transplantation Enhances Physical Function and Biological Potency in Mice. (A-C) Changes in serum levels of IL-6, TNF-α, and IL-10 in acute inflammation model mice and normal model mice one week after transplantation of two different mitochondria, with untreated groups as control. n=3. (D-F) Evaluation of kidney and liver function via serum Cr, ALT, and AST levels in acute inflammation model mice and normal model mice one week after transplantation of two different mitochondria, with untreated groups as control. n=3. (G-H) Measurement of serum MDA levels and SOD activity in acute inflammation model mice and normal model mice one week after transplantation of two different mitochondria, with untreated groups as control. n=3. (I-K) Analysis of JNK and P-JNK, p65, and P-p65 protein expression levels in liver and kidney tissues via Western blot one week after transplantation of two different mitochondria in acute inflammation model mice and normal model mice, with untreated groups as control. n=3. (L) Effects of two different mitochondria on the viability of LPS-treated BMDM cells 24 hours after transplantation, with untreated groups as control. n=3. (M) Quantitative analysis of mitochondrial ROS levels in LPS-treated BMDM cells 24 hours after transplantation of two different mitochondria, with untreated groups as control. n=3. ns = no statistical significance, * = p<0.05, ** = p<0.01, *** = p<0.001, **** = p<0.0001

Both mitochondria transplantation exhibited improved cell viability in LPS-treated BMDM (Figure 6L) and lower mitochondrial ROS production (Figure 6M). Notably, the Wild Mice group demonstrated superior alleviation of oxidative stress as well as enhanced cell viability. By co-culturing the two mitochondria with normal BMDM, the enhancement of bioenergy was also observed, and the ATP level in the Wild Mice group increased more significantly (Figure S4C). This animal experiment provided a more systematic investigation into the potential of mitochondrial transplantation than the first experiment (Extend Figures 2 and 3). Although the Young C57 group showed good therapeutic effects, the robust results observed in the Wild Mice group indicated better therapeutic outcomes for both acute inflammation in mice and LPS-treated BMDM. Furthermore, this mitochondrial exhibited a greater capability to raise antioxidation ability and improve the exercise endurance of normal mice and enhance cellular bioenergy.

## Discussion

Mitochondrial transplantation, as an emerging therapeutic strategy, has shown efficacy in various disease models, attracting increasing attention (30, 48–50). Successful mitochondrial transplantation requires multiple factors, with the source of mitochondria being a critical determinant. Currently, mitochondria are primarily derived from autologous tissues or allogeneic cultured cells (1, 6, 7, 27, 51, 52), which significantly limits the clinical application of this approach. My study offers a preliminary solution to this limitation. I demonstrated the successful internalization of mitochondria from 13 heterologous sources by three different cell germlines, underscoring the safety of cross-germline mitochondrial transplantation. Given that the genomic differences between plant and animal mitochondria are even more significant than those between different animal species, I further confirmed the safety of *Vaucheria litorea* mitochondrial transplantation. These findings indicate that mitochondrial transplantation is not restricted by phylogenetic barriers. Based on this, I propose that mitochondrial transplantation is universally applicable. The unique immunological properties of mitochondria may stem from their endosymbiotic origin(53–56), while further research is needed to substantiate this hypothesis.

Mitochondria are primarily responsible for energy production, a function that is universally conserved across species(57, 58). However, they also exhibit germline-specific functional differences, highlighting their role in adaptive evolution(59–61). While current research mainly focuses on mitochondrial transplantation as an approach to provide damaged cells with energy(24, 45), emerging evidence suggests additional benefits(17, 18, 23). My results are consistent with these findings, further demonstrating that mitochondria from different germlines can offer diverse advantages in various diseases. Specifically, mitochondria derived from MDCK cells were more effective in alleviating oxidative stress in BMDMs, MDBK-derived mitochondria exhibited superior attenuation of CCCP-induced apoptosis in AC16 cells, and Vero-derived mitochondria demonstrated greater efficiency in reducing inflammation in HepG2 cells. Although these mitochondria exhibited superior performance in specific assays, no statistically significant differences were observed across all metrics, which is both acceptable and explainable. I meticulously controlled for extraneous variables, and when overall mitochondrial function was maintained at a comparable level, the observed differential benefits are likely attributable to germline-specific mitochondrial characteristics. This observation reflects a selective and reciprocal compatibility between mitochondria and recipient cells, driven by metabolic matching to the disease status of cells. The competitive internalization assay has verified this hypothesis.

Furthermore, hybrid mitochondria were generated by fusing cells from different species to create multifunctional mitochondria with diverse germline characteristics(62). By maintaining overall mitochondrial function at comparable levels, I observed results consistent with those in the previous section. Although hybrid mitochondria exhibited advantages in specific functions, they did not demonstrate absolute superiority because of the distinction between mitochondria’s primary and secondary roles. Specifically, germline-specific functions cannot surpass energy production, which remains the mitochondria’s primary role(56). This also represents the contrast between quantity and quality. In this study, I created hybrid mitochondria from rat and mouse cardiac muscle cells, two species with relatively limited phylogenetic divergence(63, 64). Rats have 21 chromosome pairs, while mice have 20, and their genomes are connected by approximately 280 homologous blocks of sequence similarity. Ninety percent of rat genes have orthologs in mice, and the two species share about 10% of their genes. However, there are still significant differences, including genes related to immune response and protein degradation(65–69). Given this context, hybrid mitochondria derived from fused cells did not acquire sufficient germline-specific traits to surpass primary mitochondrial functions. Although the overall function of hybrid mitochondria was not markedly improved, they demonstrated superior therapeutic potential compared to single-germline mitochondria, likely due to functional optimization and selective compatibility with recipient cells.

Notably, the more powerful therapeutic outcomes could be driven by the superior functionality of mitochondria with robust overall performance. In the alleviation of HepG2 apoptosis, inflammation in BMDM cells, and HepG2 senescence, mitochondria from Bullfrog, Salmon, and PK15 cells consistently exhibited significantly greater therapeutic efficacy than other tested mitochondria. These results suggest that, at the current stage of research, mitochondria with superior overall performance are more suitable for therapeutic applications. The mitochondria of normal HepG2 cells internalized were significantly fewer than those of functionally impaired HepG2 cells (Figures S2A-C), indicating the variable quality of extracted mitochondria and differential internalization patterns based on cellular needs. This selective process resembles a “refinement” mechanism, exemplifying the bidirectional selection between mitochondria and recipient cells. The overarching biological implication of this study is that mitochondria with greater bioenergetic capacity are more conducive to survival, echoing the classic “survival of the fittest” paradigm of natural selection.

Mitochondrial transplantation has shown promising results in clinical trials for ischemic heart disease and severe late-stage diabetes mellitus (SLSDM)(1, 70). While autologous(1, 20) and allogeneic(52, 71, 72) mitochondria are preferable sources, extensive research has already demonstrated the safety of xenogeneic mitochondrial transplantation(27, 71, 73). My study further corroborates these findings and expands their applicability across a broader range of germlines. The outcomes of these clinical trials, particularly in SLSDM, could show good therapeutic effects. To investigate this, I explored the therapeutic potential of mitochondrial transplantation in animal models, based on prior in vitro experiments. Specifically, a significant therapeutic advantage was observed, with mitochondria possessing superior overall functionality, across both allogeneic and xenogeneic transplants. This suggests that mitochondria sourced from physically fit wild animals or well-conditioned individuals may serve as optimal candidates for this technique(74, 75). This insight holds particular relevance for future human studies, where autologous mitochondria from patients may be suboptimal due to disease-related impairments. In such cases, mitochondria from healthier, younger allogeneic, or even xenogeneic donors may offer enhanced functionality and improved therapeutic effects(76–79). My findings are consistent with those from mitochondrial competitive internalization assays, demonstrating that while exercise may increase oxidative stress damage(80, 81), the competition among donor mitochondria promotes the internalization of more robust mitochondria and rejects those with inferior function. This result was consistently observed in both rounds of in vivo experiments, where I recorded significant enhancements in biological function. Notably, mitochondria from wild mice exhibited superior therapeutic efficacy, suggesting that mitochondria from highly fit animals with enhanced mitochondrial functionality represent an optimal choice for transplantation. Additionally, wild mice mitochondria not only improved disease models but also enhanced the biological function of normal recipient cells, significantly expanding the potential applications and value of mitochondrial transplantation. These findings also provide a potential solution to tolerance issues that may arise with repeated mitochondrial transplantation in future therapeutic regimens.

Despite these promising findings, this study has several limitations that should be acknowledged. First, I did not evaluate mitochondrial heterogeneity, which is a crucial factor in safety assessments. Second, my research may lack proteomic and metabolomic data, which are essential for understanding the compatibility between recipient cells and transplanted mitochondria, as well as for comparing pre- and post-transplant functional and metabolic changes. Therefore, future studies should integrate these data to establish a comprehensive database of multispecies mitochondria, enabling more precise matching to specific diseases and thus providing personalized and more effective clinical solutions. Furthermore, some experiments could not be completed due to funding and other reasons constraints, highlighting the need for further research to strengthen the evidence base for future advancements in the field.

In summary, my results demonstrate that mitochondrial transplantation is not constrained by phylogenetic barriers, with mitochondria from diverse species could provide adaptable and versatile therapeutic options for patients. Moreover, my study indicates that this transplantation could serve as a bio-enhancement strategy for healthy individuals. These findings offer significant insights and lay a solid foundation for the future clinical application of mitochondrial transplantation.

## Material and methods

### Reagents and Antibodies

DMEM (C11965500BT), SF-900™ II SEM (10902088), FBS (10099141C), Penicillin-Streptomycin (15140122), and phenol red-free DMEM (12348017) were purchased from Gibco (USA). ProLong™ Anti-fade reagent (P36975) was obtained from Invitrogen (USA). Wheat Germ Agglutinin (WGA594) (W11262), Wheat Germ Agglutinin (WGA488) (W11261), Wheat Germ Agglutinin (WGA647) (W32466), MitoTracker™ Deep Red FM (M22426), MitoTracker™ Green FM (M7514), MitoTracker™ Red CMXRos (M7512), BCA Protein Assay Kit (23250), and SYBR™ Green PCR Master Mix (4309155) were purchased from ThermoFisher (USA). Lipopolysaccharide (LPS) (L2880), Hydrogen peroxide solution (H_2_O_2_) (323381), and PEG1450 solution (50% sterile-filtered) (P7181) were obtained from Sigma Aldrich (USA). D-Galactose (D-gal) (HY-N0210) and MitoSox Red (HY-D1055) were purchased from MCE (USA). Carbonylcyanide-m-chlorophenylhydrazone (CCCP) was obtained from Abcam (UK). Mercuric chloride (HgCl2) (10013616) was purchased from Sinoreagent (Shanghai, China). Mitochondria-Cytosol Protein Isolation Kit (C0010-50) was purchased from Applygen (Beijing, China), and Plant Mitochondrial Extraction Kit (SM0020) was also obtained from Beijing, China. PrimeScript™ RT Reagent Kit (RR037Q) was purchased from Takara (Japan). Mitochondrial Complex III Activity Assay Kit (E-BC-K149-M) and Mitochondrial Complex I Activity Assay Kit (E-BC-K151-M) were obtained from Elabscience (Wuhan, China). ATP Assay Kit (S0026), Mitochondrial Membrane Potential Assay Kit with JC-1 (C2006), Cell Counting Kit-8 (C0037), Mouse IL-6 ELISA Kit (PI326), Mouse IL-10 ELISA Kit (PI522), and Mouse TNF-α ELISA Kit (PT512) were purchased from Beyotime (Jiangsu, China). Superoxide Dismutase (SOD) Assay Kit (A001-2-2), Malondialdehyde (MDA) Assay Kit (A003-1-2), Glutathione Peroxidase (GSH-PX) Assay Kit (A005-1-2), Total Nitric Oxide Synthase Assay Kit (A014-2-2), Aspartate Aminotransferase Assay Kit (C010-2-1), Alanine Aminotransferase Assay Kit (C009-2-1), and Creatinine (Cr) Assay Kit (C011-2-1) were obtained from NJJCBIO (Nanjing, China).

The following antibodies were used for Western blotting: anti-p65 (1:1000, Abcam, ab32536), anti-phospho-p65 (1:1000, Cell Signaling, 3033), anti-JNK (1:1000, Cell Signaling, 9252), anti-phospho-JNK (1:1000, Cell Signaling, 9251), anti-Drp1 (1:1000, Cell Signaling, 8570), anti-Mfn1 (1:1000, Cell Signaling, 14739), anti-Opa1 (1:1000, Cell Signaling, 80471), β-actin (1:1000, Cell Signaling, 4970), and Goat Anti-Rabbit IgG (H+L) (1:3000, Abmart, M213811S).

### Cell Lines

Bovine kidney cells (MDBK) (GNO7), porcine kidney cells (PK15) (GNO31), African green monkey kidney cells (Vero) (SCSP-520), feline kidney cells (CRFK) (GNO16), canine kidney cells (MDCK) (GNO23), human cardiomyocytes (AC16) (SCSP-555), rat cardiomyocytes (H9C2) (GNR5), mouse cardiomyocytes (HL1) (GNM46), mouse myoblasts (C2C12) (SCSP-505), and human hepatocellular carcinoma cells (HepG2) (SCSP-510) were obtained from the Cell Resource Center, Chinese Academy of Sciences (Beijing, China). Chicken hepatocellular carcinoma cells (LMH) (SNL-494) were purchased from Sunncell (Wuhan, China), while Spodoptera frugiperda cells (Sf9) (CL-0205) and mouse fibroblasts (L929) (CL-0089) were obtained from Procell (Wuhan, China). MDCK, MDBK, Vero, PK15, CRFK, LMH, AC16, H9C2, HL1, C2C12, and L929, HepG2 were cultured in DMEM (Gibco) supplemented with 10% (v/v) FBS (Gibco) and 1% (v/v) penicillin-streptomycin (Gibco) at 37°C in a humidified incubator with 5% CO₂. Sf9 were cultured in SF-900™ II SEM (Gibco) medium under suspension conditions in a shaker flask at 28°C.

### Animals

The animal experiments conducted in this study utilized C57BL/6J strains, which were all C57 B/L 6J male mice purchased from the Animal Protection Center of Southern Medical University. All mice were maintained in a specific pathogen-free SPF environment. All mice were used in accordance with the scheme approved by the Review Committee of Southern Medical University. The mouse experimental scheme has also been approved by the Animal Care and Use Committee of Southern Medical University, and the experimental animal facilities are licensed by the Guangdong Medical Experimental Animal Center (SCXK-2021-0041). Additionally, lizards, salmon, bullfrogs, sparrows, turtles, and eels were acquired from animal breeding markets. Wild mice were captured using traps in their natural habitat, disinfected, and subsequently housed in the laboratory under appropriate conditions.

### Human Subjects

The human subject was the author (Xiaomeng Lu, sub-healthy), who provided full informed consent. Muscle tissue was obtained from the gastrocnemius muscle following standard surgical procedures on September 30, 2024. All procedures were conducted in strict compliance with the ethical guidelines approved by the Institutional Review Board of Southern Medical University.

### Plants

Vaucheria litorea was collected and transported under expert supervision and used immediately after thorough disinfection.

## Detailed Methods

### Isolation of Mouse Bone Marrow-Derived Macrophages (BMDMs)

Bones were aseptically harvested from the hind legs of C57BL/6 mice, and muscle tissues were removed. Bone marrow was flushed out of the bones using a 25-gauge needle attached to a syringe containing BMDM growth medium, which consists of DMEM (Gibco), 20% L929 cell-conditioned medium to generate M-CSF, 10% FBS (Gibco), and 1% penicillin/streptomycin (Gibco). BMDMs were allowed to differentiate for 7 days at 37°C with 5% CO2, with the growth medium changed every 2 days during ex vivo culture.

### Establishment of Cell Disease Models

Oxidative stress models were created by treating both BMDM and AC16 cells with 500 µM H2O2 (Sigma) for 24 hours and HepG2 cells with 700 µM H2O2 (Sigma) for 24 hours. Inflammation models were developed by treating BMDM cells with 100 nM LPS (Sigma) for 24 hours and HepG2 cells with 1 µM LPS (Sigma) for 24 hours. Mitophagy models were established by treating both AC16 and HepG2 cells with 30 µM CCCP (Abcam) for 24 hours. The senescence model was created by treating HepG2 cells with 30 g/L D-gal (MCE) for 48 hours.

### Disinfection and Sterilization of Plant Mitochondria

Weigh 1 g of tissue and rinse it with clean water for 3 hours. Transfer the tissue to a laminar flow hood and soak it in 75% ethanol for 1 minute. Then, wash the tissue twice with DPBS containing 1% streptomycin/penicillin (Gibco). After soaking in 2% NaClO for 8 minutes, wash the tissue three times with DPBS containing 1% streptomycin/penicillin (Gibco), rinsing for 1 minute each time. Next, rinse the tissue repeatedly in a 1% mercuric chloride (Sinoreagent) solution for 10 minutes. Afterward, wash the tissue three more times with DPBS containing 1% streptomycin/penicillin (Gibco). Finally, place the sterilized tissue blocks into a 100 mm² culture dish (Corning, USA, 43107) and cut them into 0.1 cm² pieces for further use.

### Mitochondrial Isolation

Mitochondria were isolated from cultured cells, animal tissues, and plant tissues using the Mitochondria-Cytosol Protein Isolation Kit (Applygen) and the Plant Mitochondrial Extraction Kit (Solarbio). For tissue homogenization, 100–200 mg of fresh animal tissue or 1 g of plant tissue was cut into 0.5 cm² pieces and placed into a Teflon-glass homogenizer containing 2.5 ml of ice-cold mitochondrial extraction buffer. The tissue was homogenized 20 times using a tight-fitting pestle. For cell homogenization, 2-5 × 10⁷ cells were collected, digested, centrifuged, washed, and resuspended in 1.5 ml of ice-cold mitochondrial extraction buffer. The cell suspension was transferred to a small glass homogenizer and homogenized 30 times with a tight-fitting pestle. The homogenate was then transferred to a centrifuge tube and centrifuged at 800 × g for 5 minutes at 4°C. The supernatant was collected and transferred to a new centrifuge tube, followed by centrifugation at 10000 × g for 10 minutes at 4°C. The mitochondrial pellet was resuspended in 0.2 ml of mitochondrial extraction buffer and subjected to centrifugation at 12000 × g for 10 minutes at 4°C. The supernatant was discarded, and the mitochondrial pellet was resuspended in mitochondrial respiration buffer for subsequent use.

### Mitochondrial Quality Control

The extracted mitochondria were quantified using a BCA Protein Assay Kit (Thermo Fisher). The integrity of the isolated mitochondria was assessed with the Mitochondrial Complex I and III Activity Assay Kit (Elabscience). The mitochondrial membrane potential was measured using the Mitochondrial Membrane Potential Assay Kit with JC-1 (Beyotime), and the capacity for energy synthesis was evaluated using an ATP Assay Kit (Beyotime).

### Electron Microscopy

For transmission electron microscopy (TEM), the isolated mitochondria were fixed overnight in 0.2 M sodium cacodylate buffer containing 2.5% formaldehyde, 5% glutaraldehyde, and 0.06% picric acid at pH 7.4. The samples were washed in cacodylate buffer, stained with 1% osmium tetroxide and 1.5% potassium ferricyanide, and then dehydrated using ethanol and propylene oxide. After infiltration and embedding in Epon-Araldite (Electron Microscopy Sciences, Hatfield, PA), sections (60-80 nm thick) were cut using an Ultracut-S ultramicrotome (Reichert Technologies, Depew, NY) and mounted on copper grids (200 mesh) for observation with a transmission electron microscope (Hitachi H-7500, Japan).

### Mitochondrial Transplantation in Cells

After isolating the mitochondria, their concentration was quantified using a BCA Protein Assay Kit (Thermo Fisher). For the co-culture, 10 µg of mitochondria were added per 10⁵ cells(17). For competitive mitochondrial internalization, 1 µg of mitochondria was added per 10⁵ cells, using 0.5 µg of each type for dual mitochondrial competition or 0.33 µg of each type for triple mitochondrial competition.

### Mitochondrial Internalization and Observation

Recipient cell membranes were labeled with WGA594 (ThermoFisher) or WGA488 (ThermoFisher), while isolated exogenous mitochondria were labeled with MitoTracker™ Green FM (ThermoFisher) or MitoTracker™ Red CMXRos (ThermoFisher). After 30 minutes, both the cells and mitochondria were washed twice with DPBS. The labeled mitochondria and recipient cells were co-cultured in a 35-mm confocal culture dish (Beyotime, Jiangsu, China, FCFC020-150pcs) for 24 hours. Following the removal of the culture medium, the cells were washed twice with DPBS and fixed with glutaraldehyde. After an additional 30 minutes, the cells were washed twice with DPBS, and an antifade reagent (Prosperich, Israel, PL739) was added. Mitochondrial and cellular colocalization was observed using an inverted laser confocal microscope (LSM900 with Airyscan2, USA).

### ATP Content Measurement

ATP content was measured using an ATP Assay Kit (Beyotime) according to the manufacturer’s instructions. One hundred micrograms of isolated mitochondria or cells seeded in a 6-well plate (at a density of 5 × 10⁵ cells per well) were treated with 200 μL of ATP lysis buffer and then centrifuged at 12000 × g for 5 minutes at 4°C. Twenty microliters of the supernatant were mixed with 100 μL of ATP detection solution in a 96-well white solid flat-bottom polystyrene microplate (Corning, USA, 3599), and luminescence (RLU) was measured using a multifunctional microplate reader (Tecan, Switzerland).

### Mitochondrial Membrane Potential Fluorescence Quantification

The isolated mitochondria were subjected to protein quantification, followed by an assessment of mitochondrial membrane potential using the Mitochondrial Membrane Potential Assay Kit with JC-1 (Beyotime). In brief, 50 µg of mitochondria were mixed with 90 µL of JC-1 staining solution (2 µM) and transferred to a 96-well plate (Corning, USA, 3522). The mixture was thoroughly mixed, and the membrane potential was immediately measured using the BioTek Cytation5 Multifunctional Cell Imaging Microplate Reader (Agilent, USA), with an excitation wavelength of 485 nm and an emission wavelength of 590 nm.

### Measurement of Mitochondrial Reactive Oxygen Species (ROS) Fluorescence Intensity

Cells were seeded at a density of 1 × 10⁴ cells per well in a 96-well solid white flat-bottom polystyrene tissue culture-treated microplate (Corning, USA, 3599). After applying the respective treatment conditions, exogenous mitochondria were co-cultured with the cells for 24 hours. The medium was then removed, and the cells were washed twice with DPBS. A 5 µM working solution of MitoSOX Red (MCE) dye (100 µL) was added to each well, and the plate was gently agitated to ensure complete coverage of the cells. Following a 20-minute incubation, the dye was removed, and the cells were washed twice with the medium. The mitochondrial ROS fluorescence intensity was measured using a multifunctional microplate reader (Tecan, Switzerland) with an excitation wavelength of 510 nm and an emission wavelength of 580 nm.

### Measurement of Intracellular Reactive Oxygen Species

Cells were seeded at a density of 1 × 10⁵ cells per well in a 24-well plate (Corning, USA, 3524). After treating the cells under the specified conditions, exogenous mitochondria were co-cultured with the cells for 24 hours. The medium was then removed, and the cells were washed twice with DPBS. Subsequently, the cells were incubated with 10 µM DCFH-DA (Beyotime) or 5 µM MitoSOX Red (MCE) at 37°C for 30 minutes. The stained cells were washed with PBS and observed using the LSM900 with Airyscan2 (Zeiss, Germany) or the EVOS M5000 (ThermoFisher, USA).

### Measurement of Mitochondrial Membrane Potential

Cells were seeded at a density of 1 × 10⁵ cells per well in a 24-well plate (Corning, USA, 3524). After treating the cells under the specified conditions, exogenous mitochondria were co-cultured with the cells for 24 hours. The medium was subsequently removed, and the cells were washed twice with DPBS. Following the protocol for the Mitochondrial Membrane Potential Assay Kit using JC-1, 500 µL of JC-1 staining solution (2 µM) was added to each well and incubated for 20 minutes. The cells were then washed and observed using the LSM900 with Airyscan2 (Zeiss, Germany) or the EVOS M5000 (Thermo Fisher, USA) in phenol-red-free DMEM (Gibco).

### CCK8 Assay

According to the Cell Counting Kit-8 (Beyotime) protocol, 100 µL of cell suspension (density of 1 × 10⁴ cells/mL) was seeded in a 96-well plate (Corning, USA, 3522). After treating the cells under the specified conditions, 10 µL of CCK8 reagent was added, and the cells were incubated at 37°C for 2 hours. The absorbance at 450 nm was measured using a multifunctional microplate reader (Tecan, Switzerland). The percentage of cell viability was calculated based on the average optical density (OD) of each group.

### DNA Oxidative Damage Detection

The supernatant from the cultured cells was collected 24 hours after mitochondrial transplantation. The concentration of 8-OHdG was measured using the 8-OHdG ELISA Kit (Elabscience), and absorbance at 450 nm was recorded to quantify DNA oxidative damage.

### Immunoinflammatory Cytokines Detection

The supernatant from the cultured cells was collected 24 hours after mitochondrial transplantation. The concentrations of IL-6, IL-10, and TNF-α were measured using the Mouse IL-6 ELISA Kit (Beyotime), Mouse IL-10 ELISA Kit (Beyotime), and Mouse TNF-α ELISA Kit (Beyotime), respectively, following the manufacturer’s instructions, with absorbance readings taken at 450 nm.

### Measurement of MDA, GSH, SOD, ALT, AST, Cr

The supernatants of cultured cells following mitochondrial transplantation, as well as serum from model mice, were collected. According to the manufacturer’s instructions (NJJCBIO), the optical density (OD) values for MDA, GSH, SOD, ALT, AST, and Cr were measured at wavelengths of 532 nm, 412 nm, 550 nm, 505 nm, 510 nm, and 546 nm, respectively.

### Cell Fusion

When the cells reached high proliferation rates and optimal conditions, 3×10⁶ HL-1 and H9C2 cells were harvested separately. HL1 cells were stained with WGA647, while H9C2 cells were stained with WGA594. The staining was conducted at 37°C for 25 minutes, followed by two washes with DPBS. The cells were then combined and centrifuged at 200 g for 5 minutes. The supernatant was discarded, and pre-warmed 50% PEG1450 (Sigma) was gradually added while stirring the pellet. A total of 700 µL of 50% PEG1450 (Sigma) was added within 1 minute. The mixture was incubated at 37°C for 1 minute, after which 10 mL of DMEM was added to dilute the PEG1450. The centrifuge tube was placed in a 37°C incubator for 10 minutes before being centrifuged again at 200 g for 5 minutes. The cells were washed twice with DPBS, resuspended in complete culture medium, and incubated at 37°C with 5% CO₂ for 24-48 hours. Cell fusion was observed using the LSM900 with Airyscan2 (Zeiss, Germany), and fused HL1+H9C2 cells were sorted using a MoFlow XDP flow sorter (Beckman, USA) for further culture.

### Competitive Internalization of Mitochondria

For the competitive internalization of two types of mitochondria, the mitochondria were stained with MitoTracker™ Deep Red FM (Thermo Fisher) and MitoTracker™ Green FM (Thermo Fisher) for 30 minutes. Following two washes with DPBS, the mitochondria were co-cultured with recipient cells for 24 hours. The medium was then replaced with phenol red-free medium containing ProLong™ antifade reagent (Invitrogen), and live-cell imaging was conducted using the LSM900 with Airyscan2 (Zeiss, Germany). For the competitive internalization of three types of mitochondria, the mitochondria were stained with MitoTracker™ Deep Red FM (Thermo Fisher), MitoTracker™ Green FM (Thermo Fisher), and MitoTracker ™ Red CMXRos (Thermo Fisher) for 30 minutes. After two washes with DPBS, the mitochondria were co-cultured with recipient cells for 24 hours. The medium was then replaced with phenol red-free DMEM (Gibco) containing ProLong™ antifade reagent (Invitrogen), and live-cell imaging was performed using the LSM900 with Airyscan2 (Zeiss, Germany).

### HIIT Functional Training

Three 3-month-old mice underwent a one-week pre-adaptation exercise program on a small animal treadmill (Shxinruan) before commencing an 8-week high-intensity interval training (HIIT) regimen. The maximum exercise capacity was assessed weekly to adjust the running speed accordingly. Each session began with a 10-minute warm-up at a speed of 10 m/min. During the first two weeks, the regimen included high-intensity exercise at 24 m/min (80% VO2max) for 3 minutes, followed by 3 minutes of low-intensity recovery at 15 m/min with a 0° incline. In weeks 3 and 4, the regimen increased to 26 m/min (80% VO2max) for 3 minutes, followed by 3 minutes at 15 m/min with an 8° incline. In weeks 5 and 6, the speed was further increased to 28 m/min (80% VO2max) for 3 minutes, followed by 3 minutes at 17 m/min with a 16° incline. Finally, in weeks 7 and 8, the speed reached 30 m/min (80% VO2max) for 3 minutes, followed by 3 minutes at 17 m/min with a 25° incline. Each session concluded with 8 minutes at 15 m/min. The regimen consisted of 7 sets (3+3) per session, with each session lasting 1 hour, conducted 5 days per week, and included 2 days of rest. Similarly, a 30-year-old sub-healthy human (Xiaomeng Lu) participated in an 8-week HIIT training program, which involved a 30-second maximal effort sprint (rated 9+ on the 1–10 RPE scale) followed by a 4.5-minute low-intensity jog (rated 4–5 on the 1–10 RPE scale), performed daily for 30 minutes, 5 days per week, along with an additional 2 days of moderate-intensity continuous training (MICT) running at 15 km/h for 30 minutes.

### Aging Model

The aging model was categorized into two categories. Disease Model: A total of 45 three-month-old C57 B/L mice were randomly assigned to five groups: D-gal group, C2C12 group, Young C57 group, Trained People group, Trained C57 group, and Wild Mice group, with 9 mice in each group. Aging was induced through subcutaneous injection of D-gal (MCE) at a dosage of 500 mg/kg once daily for 8 consecutive weeks. Normal Model: A total of 27 five-month-old C57 B/L mice were divided into three groups: Ctrl group, Young C57 group, and Wild Mice group, with 9 mice in each group. These mice received no drug treatment and were maintained under standard conditions.

### Acute Inflammation Model

The purchased two-month-old mice were not treated and normally cultured to four-month-old. The acute inflammation model was categorized into two categories. A total of 27 four-month-old C57BL/6 mice were divided into three groups: the LPS group, the Young C57 group, and the Wild Mice group, with 9 mice in each group. Acute inflammation was induced through intraperitoneal injection of LPS (Sigma) at a dosage of 2.5 mg/kg/day for 5 consecutive days. Normal Model: Additionally, another 27 four-month-old C57BL/6 mice were divided into three groups: the Ctrl group, the Young C57 group, and the Wild Mice group, with 9 mice in each group. These mice received no drug treatment and were maintained under standard conditions.

### Animal Mitochondrial Transplantation

For the aging model, mitochondria were isolated from the gastrocnemius muscle tissues of five-month-old Young C57, Trained C57, Trained People and Wild Mice, as well as C2C12 cells, using the Mitochondria-Cytosol Protein Isolation Kit (Applygen). Mitochondrial protein concentration was quantified using the BCA Protein Assay Kit (ThermoFisher). The isolated mitochondria were resuspended in DPBS and injected via the tail vein at a dosage of 100 μL/100 μg(82) into mice from the C2C12 group, Young C57 group, Trained C57 group, Trained People group and Wild Mice group, while the D-gal group received no treatment. For the normal model, mitochondria from Trained C57 and Wild Mice were injected into their respective groups at the same dosage, with the Ctrl group receiving 100 μL of mitochondrial carrier solution. In the acute inflammation model, mitochondria were isolated from the gastrocnemius muscle of two-month-old Young C57 and Wild Mice, quantified, and injected via the tail vein at a dosage of 100 μL/100 μg into the Young C57 and Wild Mice groups. The LPS group received no treatment. For normal model, Young C57 group and Wild Mice group were injected 100 μL/100 μg with the corresponding mitochondria through the tail vein, and the Ctrl group received 100 μL of mitochondrial carrier solution.

### Mitochondrial Biodistribution

The isolated mitochondria were stained with MitoTracker™ Deep-Red FM (ThermoFisher) at room temperature for 20 minutes, followed by two washes with DPBS. The mitochondria were then resuspended in DPBS and injected into mice via the tail vein. After 24 hours, the hearts, livers, and kidneys were harvested, and the in vivo biodistribution of the mitochondria was analyzed using the PerkinElmer IVIS Spectrum (PerkinElmer, USA) multimodal small animal imaging system.

### Motor Function Detection (Rotarod Test)

Adaptive training was performed before the experiment. During the formal experiment, the mice were placed on a rotating rod apparatus (Shxinruan, China) set to 5 rpm/min for 5 minutes, after which the speed was gradually increased to 40 rpm/min. The duration for which each mouse remained on the rotating rod was recorded using a stopwatch until they fell off. The maximum duration allowed for aging or acutely inflamed mice was 180 seconds, while normal mice were permitted a maximum of 360 seconds. Each mouse was tested three times, with a 3-minute interval between trials, ensuring a quiet environment throughout the experiment.

### Body Weight Measurement

All experimental groups of mice were weighed three days before and one week after mitochondrial transplantation, and their weights were recorded.

### Mouse Forelimb Grip Strength Test

Adaptive training was performed before the experiment. During the formal experiment, each mouse was positioned on a mouse grip tester (Shxinruan, China), and gentle traction was applied to the tail, encouraging the mouse to grip the probe firmly with its forelimbs. The grip strength reading was recorded when the experimenter applied maximum force, and the experiment was conducted in a quiet environment.

### Lower Limb Muscle Volume and Injury Assessment

One week after mitochondrial injection, mice were briefly anesthetized with a mixture of isoflurane (3.0%) and O2 (2.0 L/min) via a nose cone and placed in a custom-made restraint tube, with the right lower limb fixed with a coil. During MRI imaging, mice were anesthetized with a mixture of isoflurane (1.5%) and O2 (0.6 L/min). The rectal temperature was maintained at 37 ± 1°C using a heated water blanket. A small animal magnetic resonance imaging system, PharmaScan70/16S (Bruker, USA), was used to scan the horizontal plane of lower limb muscles, employing spin echo (SE) T1WI (TR4000ms, TE 28.5ms), fast spin echo (FSE) T2WI (TR4000ms, TE 47.5ms), with a slice thickness of 4 mm and an interval of 1 mm. Process the images using Paravision6.0.1 (Bruker Software, USA).

### Western Blot

For Cells: On the second day after cell fusion, HL1, H9C2, and HL1+H9C2 cells were lysed, and proteins were extracted from each cell type. For Tissues: Mice were randomly selected from each of the two experimental groups in the second animal experiment. Three mice were chosen from each group of the acute inflammation model and three from each group of the normal model. Proteins were then extracted from the liver and kidney tissues of these mice. Protein concentrations were measured using a BCA protein assay kit. Proteins were separated by 10% sodium dodecyl sulfate-polyacrylamide gel electrophoresis (SDS-PAGE) and subsequently transferred onto polyvinylidene fluoride (PVDF) (Millipore, USA) membranes. The membranes were blocked with 5% bovine serum albumin (BSA) or 5% milk for detecting phosphorylated proteins (P-p65, P-JNK) and total proteins (p65, JNK, β-actin, Drp1, Mfn1, Opa1). The PVDF membranes were incubated with primary antibodies at 4°C overnight, followed by incubation with horseradish peroxidase-conjugated secondary antibodies at room temperature for 2 hours. Gel images were captured using the Automatic Luminous System Tanon-5200Multi (Tanon, China), and data were analyzed using ImageJ (ImageJ Software, National Institutes of Health, USA).

### Quantitative PCR

Twenty-four hours after mitochondrial transplantation, total RNA was extracted using Trizol reagent. Purified RNA was reverse-transcribed into cDNA using the PrimeScript™ RT reagent Kit (Takara). RT-qPCR was conducted with SYBR Green Master Mix (ThermoFisher) using the ABI QuantStudio 5 Real-Time PCR system (ThermoFisher, USA). Expression levels were normalized to the internal control 18s rRNA. The primer sets are listed in Key resources table.

### Quantification and Statistical Analysis

Cell experiments were repeated at least three times except for mitochondrial competitive internalization and animal experiments except for body weight measurement, grip strength and motor function tests. GraphPad Prism 9.01 (GraphPad Software Inc., USA) software was used for statistical analysis. The data were expressed as mean ± standard error (SEM). For normally distributed data and comparisons between two groups, the significance was calculated using unpaired two-tailed Student’s *t*-tests. Comparisons between three or more groups were performed using ordinary One-way ANOVA, Using Two-way ANOVA and Bonferroni post hoc test were used to compare multi-factor variance for multiple data. The significance threshold was determined by p value < 0.05, and the annotation was ns = no statistical significance, * = p<0.05, ** = p<0.01, *** = p<0.001, **** = p<0.0001.

## ACKNOWLEDGMENTS

I am very grateful to Professor Yong Jiang of Southern Medical University for the experimental site and many equipment support provided for this experiment. I sincerely thank the People’s Hospital of Dongguan City, affiliated with Southern Medical University, for providing free access to experimental instruments. I also extend our gratitude to Dr. Guiming Chen from the Department of Pathology and Pathophysiology at Southern Medical University and Dr. Rong Wu from the School of Traditional Chinese Medicine for their technical support with mitochondrial tail vein injections. Additionally, I appreciate the assistance of Master Xin Heng in the experiments involving fluorescence microscopy for detecting reactive oxygen species and mitochondrial membrane potential. Moreover, I express strong condemnation towards those who deliberately sabotage others’ experiments. I call for all laboratories actively weed out the bad apples.

## Author Contribution

Xiaomeng Lu leads all the experimental design, experimental operation, experimental analysis, article writing and proofreading.

## Funding

No founding, this work was carried out with the Xiaomeng Lu’s full commitment, with all funding provided personally by the author(Xiaomeng Lu).

### Availability of data and materials

All data generated in the present study may be requested from the corresponding author.

## Declarations

### Consent for publication

The author declares no competing interests.

### Ethics approval and consent to participate

The experimental animal facilities have been licensed by the Guangdong Medical Experimental Animal Center (SCXK-2021-0041).

### Consent for publication

Not applicable.

### Contributor Information

Xiaomeng Lu.

---

**Figure S1.**
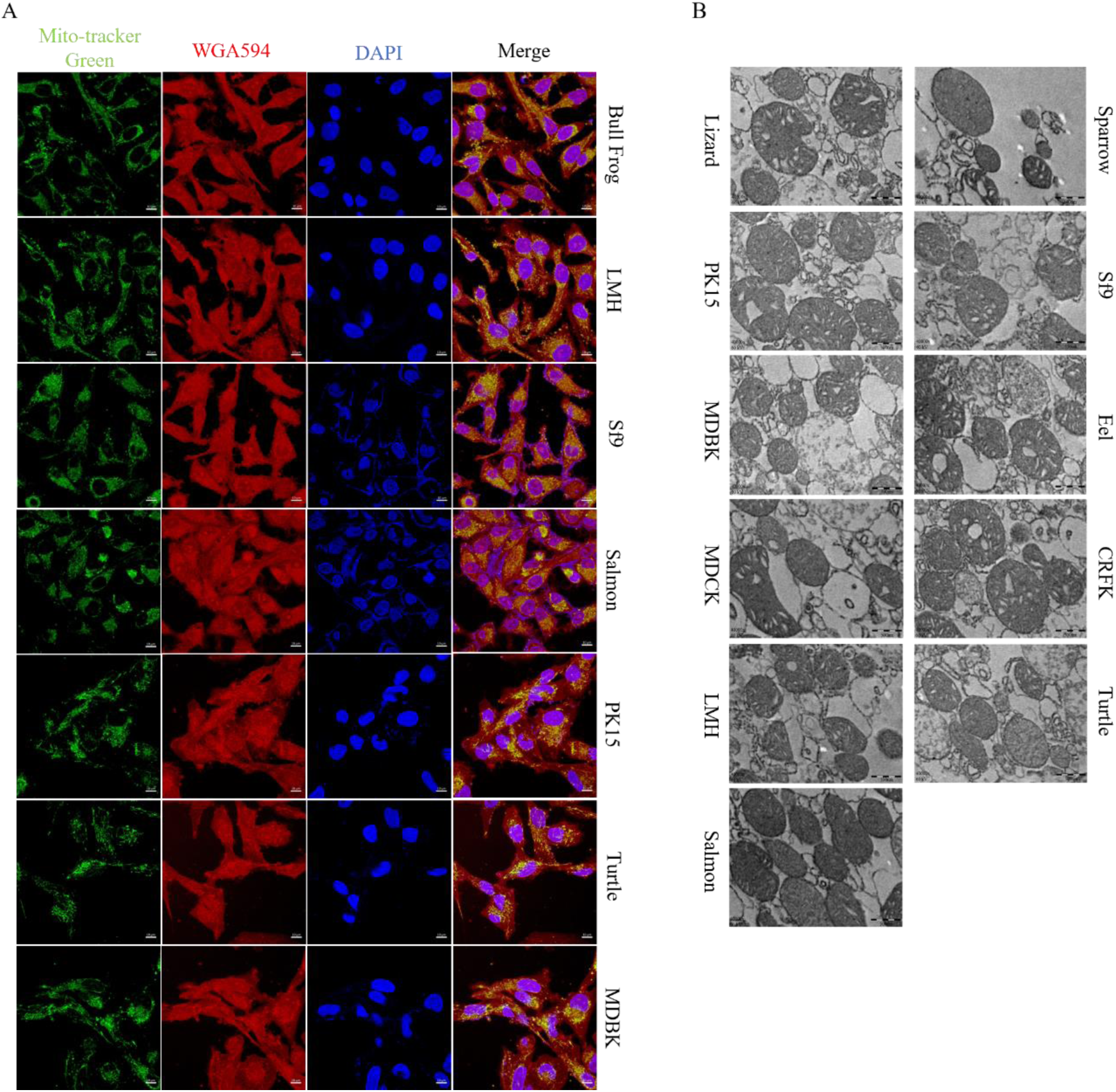
Data related to Figure 1. (A) Mitochondria labeled with green fluorescence using MitoTracker Green were transplanted from Salmon, Sf9, PK15, LMH, Turtle, MDBK and Bull Frog, and their colocalization with red fluorescence-labeled AC16, L929, and HepG2 cells stained with WGA594 was observed using fluorescence imaging after 24 hours. Blue fluorescence (DAPI) represents the cell nuclei. Scale bar: 10 µm. (B) Mitochondria from cells (MDBK, MDCK, CRFK, PK15, Sf9, LMH) and animals (Lizard, Sparrow, Salmon, Eel, Turtle) for electron microscopy to observe their morphology. Scale bar: 500 nm.

**Figure S2.**
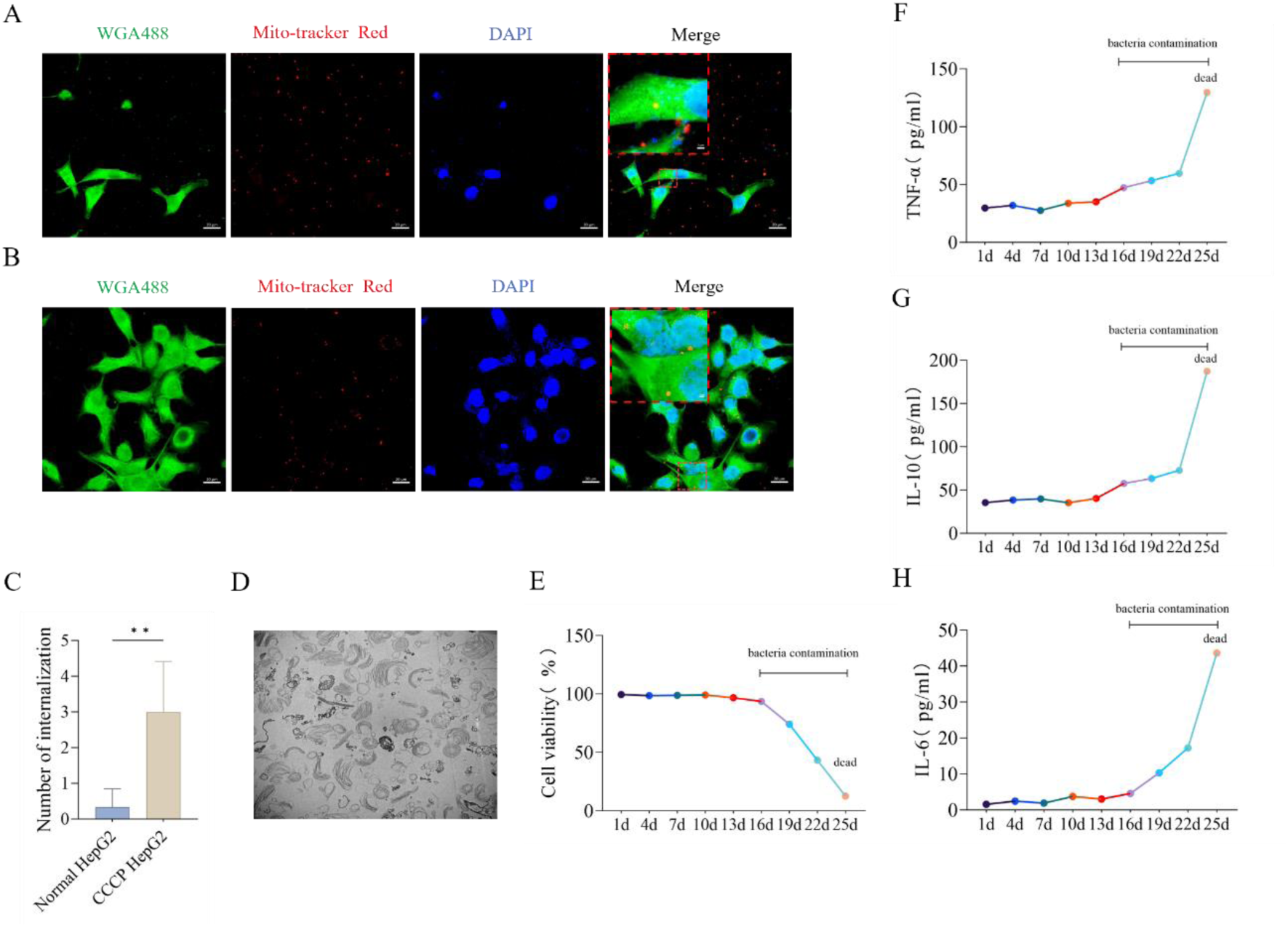
Data related to Figure 1 and Figure 4. (A) Co-localization fluorescence imaging of CCCP-treated HepG2 cells labeled with WGA488 and plant mitochondria labeled with Mito-Tracker Red, scale bar: 20 µm, magnified region scale bar: 2 μm. (B) Co-localization fluorescence imaging of normal HepG2 cells labeled with WGA488 and plant mitochondria labeled with Mito-Tracker Red, scale bar: 20 µm, magnified region scale bar: 2 μm. (C) Statistics of the number of Vaucheria litorea mitochondria internalized by normal HepG2 cells and CCCP-treated HepG2 cells. n=6. (D) Morphology of plant mitochondria observed under TEM, scale bar: 1 µm. (E) Changes in HepG2 cell viability after nine consecutive transplants of plant mitochondria, with untreated groups as control. (F-H) Changes in immune-inflammatory factors IL-6, IL-10, and TNF-α in HepG2 cells after nine consecutive transplants of plant mitochondria, with untreated groups as control. **: p<0.01.

**Figure S3.**
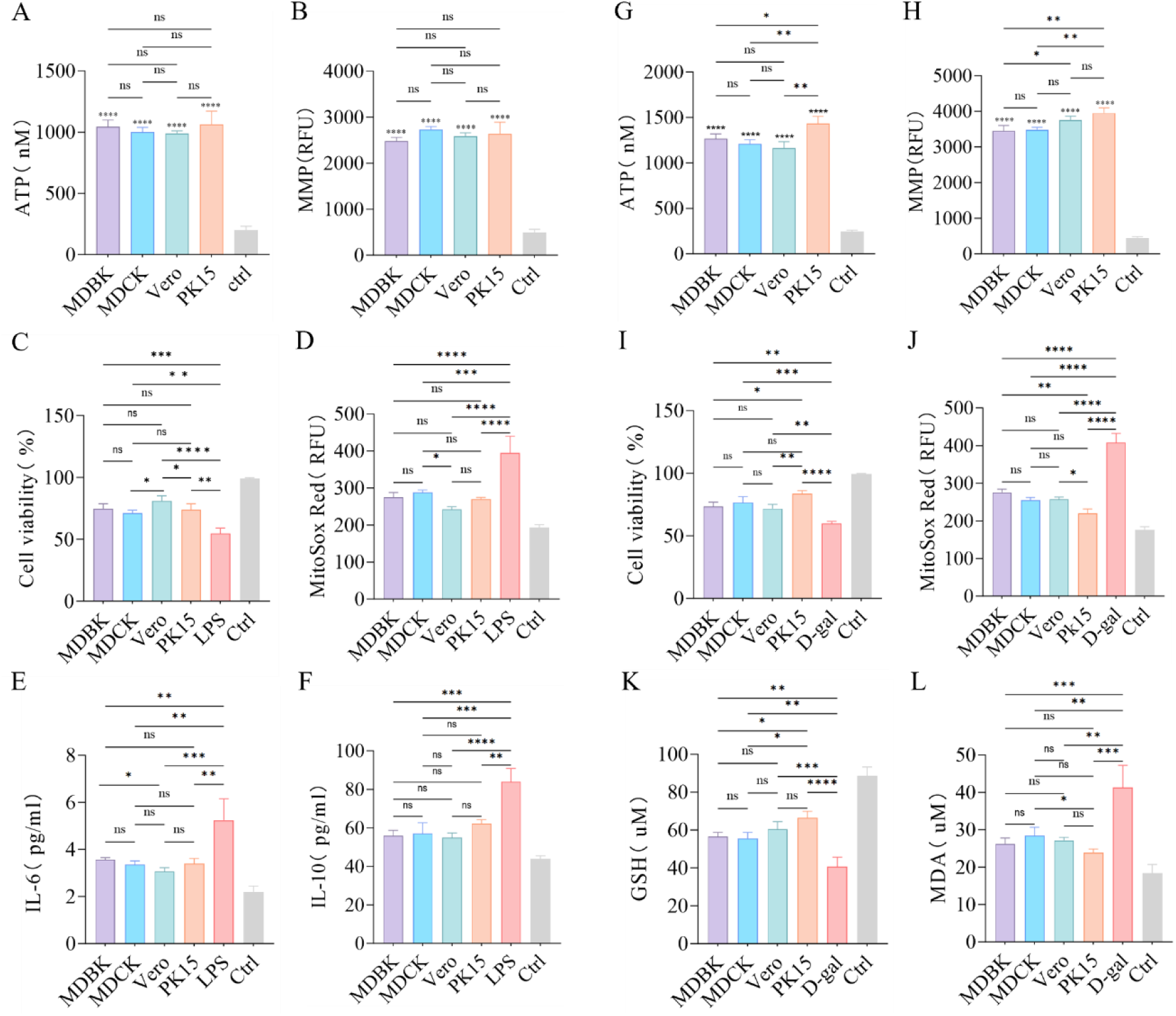
Data related to Figure 2 and Figure 4. (A, G) Assessment of ATP production capacity of isolated mitochondria, with mitochondria subjected to two freeze-thaw cycles at -80°C and 37°C to completely disrupt structure and function as control. n=3. (B, H) Analysis of mitochondrial membrane potential of isolated mitochondria, with mitochondria subjected to two freeze-thaw cycles at -80°C and 37°C to completely disrupt structure and function as control. n=3. (C) Changes in HepG2 cell viability after LPS treatment following transplantation of four different mitochondria, with untreated groups as control. n=3. (D) Effects of transplantation of four different mitochondria on mitochondrial ROS levels in LPS-treated HepG2 cells, with untreated groups as control. n=3. (E-F) Changes in IL-6 and IL-10 levels in HepG2 cells treated with LPS after transplantation of four different mitochondria, with untreated groups as control. n=3. (I) Changes in HepG2 cell viability after D-gal treatment following transplantation of four different mitochondria, with untreated groups as control. n=3. (J) Effects of transplantation of four different mitochondria on mitochndrial ROS levels in D-gal treated HepG2 cells, with untreated groups as control. n=3. (K-L) Changes in MDA and GSH levels in HepG2 cells treated with D-gal after transplantation of four different mitochondria, with untreated groups as control. n=3. ns: no statistical significance, *: p<0.05, **: p<0.01, ***: p<0.001, ****: p<0.0001. I extracted mitochondria from four types of cells and co-cultured them with LPS-treated HepG2 cells. When the overall mitochondrial functions were comparable, the Vero group exhibited superior therapeutic effects (Figures S3A-B). The Vero mitochondria significantly improved cell viability compared to the MDCK and PK15 groups (Figure S3C), reduced mitochondrial ROS production more effectively than the MDCK group (Figure S3D), and decreased IL-6 levels more substantially than the MDBK group (Figure S3E). Although there were no significant differences in IL-10 levels among the groups, the Vero group still demonstrated a slightly stronger effect (Figure S3F). These results indicate that Vero mitochondria provide better therapeutic outcomes for LPS-treated HepG2 cells, highlighting a more suitable and efficient match under these conditions. I cultured P5 generation PK15, P15 generation MDBK, P15 generation MDCK, and P15 generation Vero cells to create mitochondria with varying bioenergetic strengths, which were then co-cultured with D-gal-treated HepG2 cells. Assessments of ATP and MMP assessments revealed that mitochondria from early-passage PK15 cells exhibited stronger bioenergetic function compared to those from MDBK and Vero cells (Figures S3G-H). PK15 mitochondria significantly enhanced HepG2 cell viability (Figure S3I) and reduced ROS production (Figure S3J) in comparison to MDBK and MDCK groups. Additionally, they increased GSH levels (Figure S3K) and decreased MDA production more effectively than MDCK mitochondria (Figure S3L), thereby reducing oxidative stress in HepG2 cells. This further confirms the therapeutic advantage of mitochondria with more powerful bioenergetic function.

**Figure S4.**
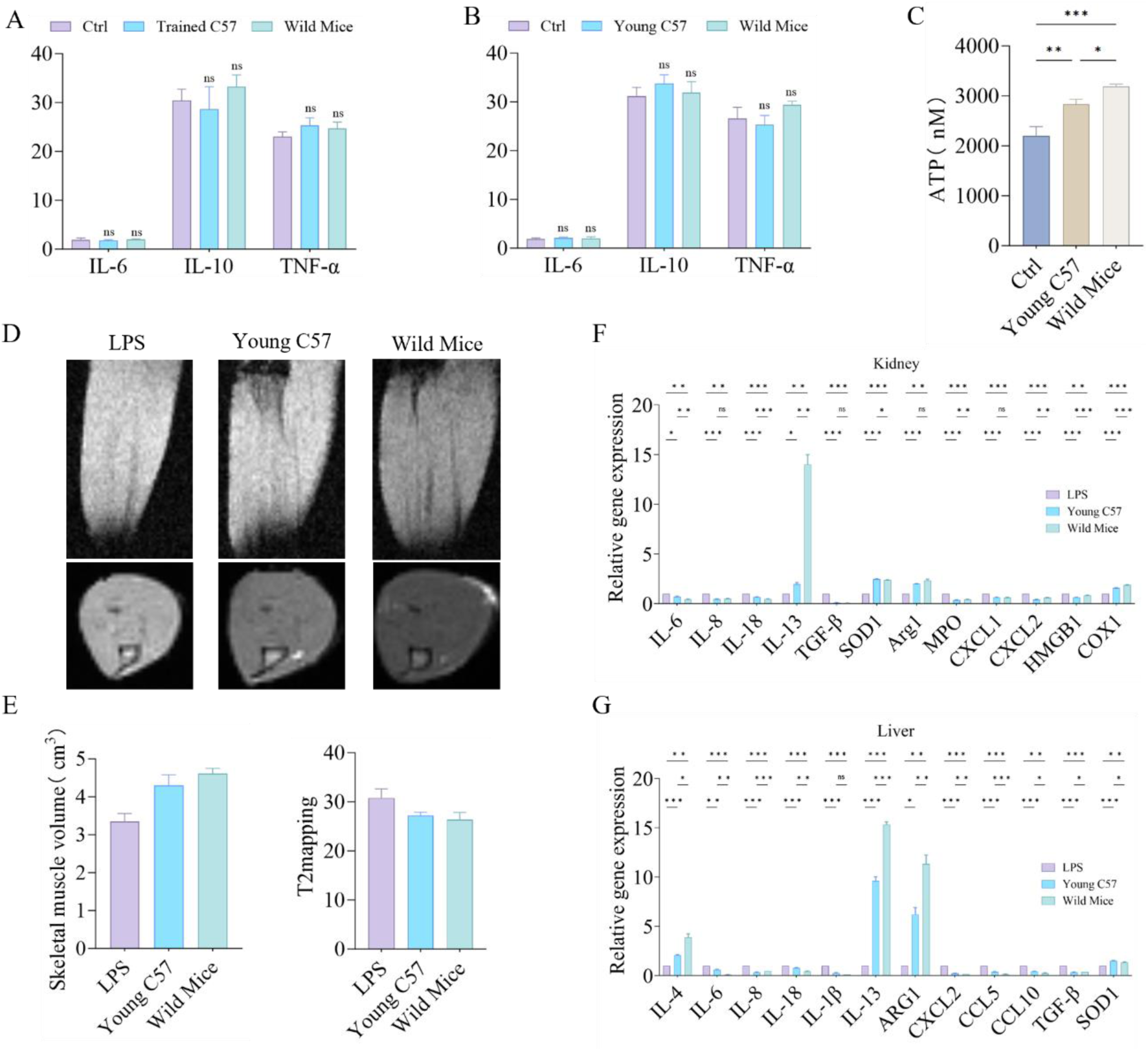
Data related to Figure 5 and Figure 6. (A) Analysis of immune-inflammatory factors IL-6, IL-10, and TNF-α in normal mice 24 hours after mitochondrial transplantation from the first animal experiment. n=3. (B) Analysis of immune-inflammatory factors IL-6, IL-10, and TNF-α in normal mice 24 hours after mitochondrial transplantation from the second animal experiment. n=3. (C) Changes in normal BMDM ATP productivity after transplantation of two different mitochondria in the second animal experiment, with untreated groups as control. n=3. (D-E) MRI assessment of hindlimb muscle volume and muscle damage repair in acute inflammation model mice one week after transplantation of different mitochondria in the second animal experiment. n=2. (F) Real-time PCR analysis of renal expression of IL-6, IL-8, IL-18, IL-13, TGF-β, SOD1, ARG1, MPO, CXCL1, CXCL2, HMGB1, and COX1 in acute inflammation model mice after transplantation of different mitochondria in the second animal experiment. n=5. (G) Real-time PCR analysis of hepatic expression of IL-4, IL-6, IL-8, IL-18, IL-1β, IL-13, TGF-β, SOD1, ARG1, CCL5, CCL10, CXCL2, and HMGB1 in acute inflammation model mice after transplantation of different mitochondria in the second animal experiment. n=5. ns: no statistical significance, *: p<0.05, **: p<0.01, ***: p<0.001, ****: p<0.0001

**Extend Figure1:**
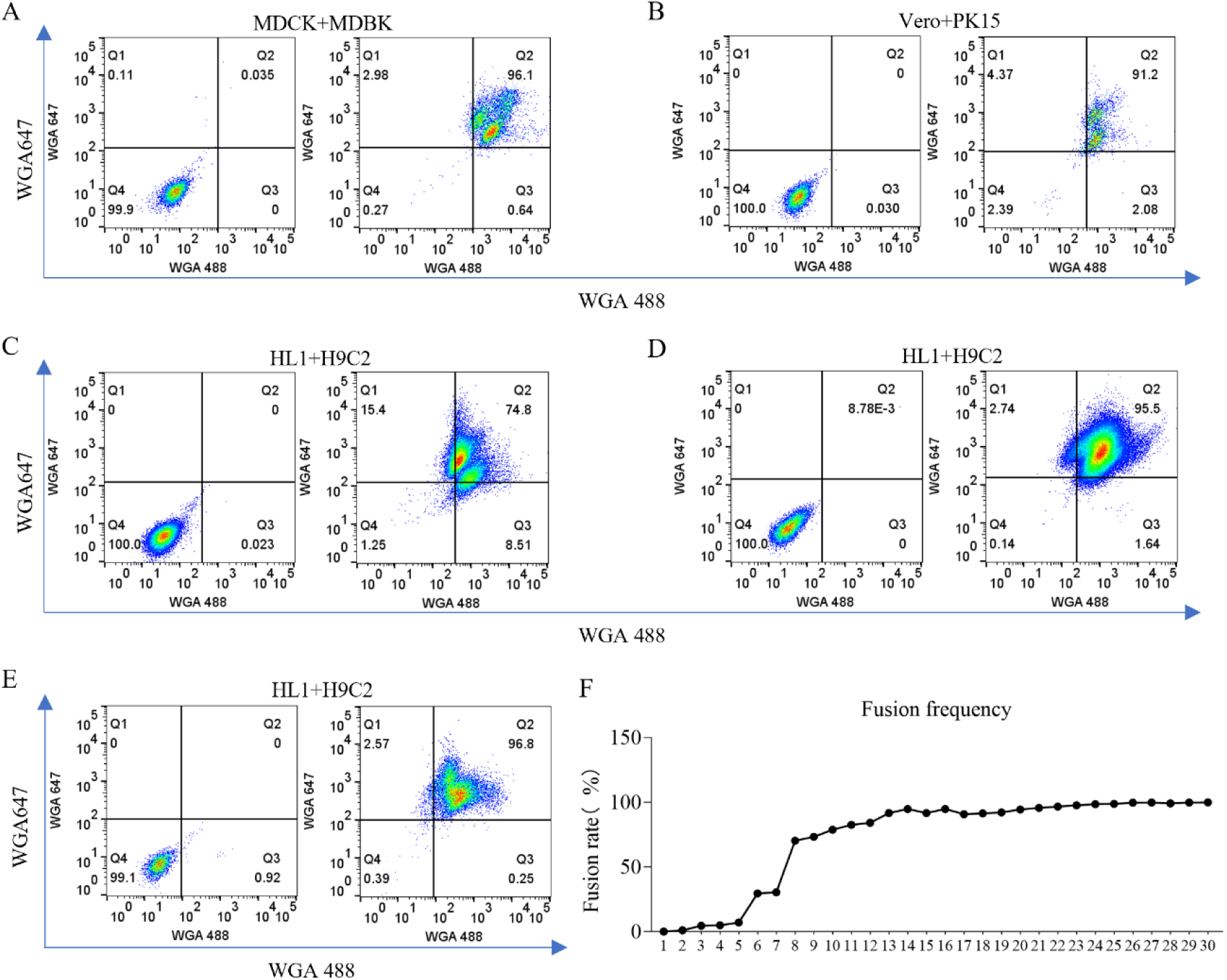
Statistical analysis of cell fusion. (A) Fusion analysis of WGA647-labeled MDBK cells and WGA488-labeled MDCK cells using flow cytometry. (B) Fusion analysis of WGA647-labeled Vero cells and WGA488-labeled PK15 cells using flow cytometry. (C-E) Fusion analysis of WGA647-labeled HL1 cells and WGA488-labeled H9C2 cells using flow cytometry. (D) Changes in cell fusion rate with improvements in experimental techniques.

**Extend Figure2 :**
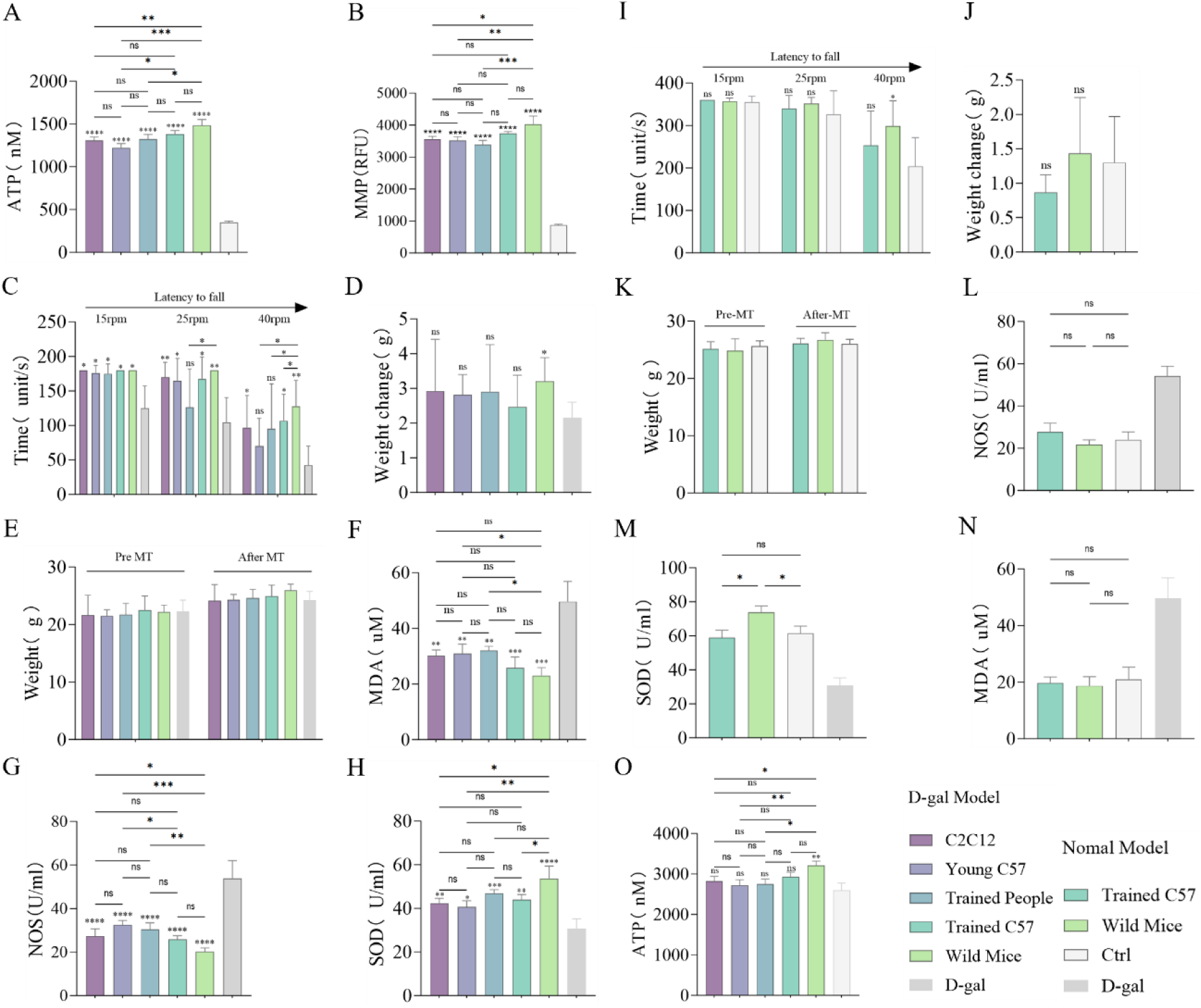
Mitochondrial Transplantation Enhances Physical Function and Biological Potency in Mice. (A-B) Measurement of ATP production capacity and membrane potential in isolated mitochondria. Ctrl: Mitochondria subjected to two freeze-thaw cycles at -80°C and 37°C, used for comparison. n=3. (C) Changes in motor ability of D-gal model mice following transplantation of five different types of mitochondria. n=8. (D-E) Changes in body weight of D-gal model mice following transplantation of five different types of mitochondria. n=8. (F-H) Evaluation of the antioxidant and oxidative capacity of D-gal model mice by measuring serum glutathione content, superoxide dismutase activity, and malondialdehyde levels after transplantation of five different types of mitochondria. n=3. (I) Changes in motor ability of normal model mice following transplantation of five different types of mitochondria. n=6. (J-K) Effects of transplantation of two different types of mitochondria on the motor ability of normal mice. Ctrl: Untreated group. n=6. (L-N) Evaluation of the antioxidant and oxidative capacity of normal mice by measuring serum GSH content, SOD activity, and MDA levels after transplantation of two different types of mitochondria. Ctrl: Untreated group. n=3. (O) Changes in ATP production capacity in normal HepG2 cells after transplantation of five different types of mitochondria. Ctrl: Untreated group. n=3. ns = no statistical significance, * = p<0.05, ** = p<0.01, *** = p<0.001, **** = p<0.0001 In the first animal experiment, I utilized a D-gal-induced mouse aging model and intravenously injected mitochondria extracted from mouse myoblasts (C2C12), gastrocnemius tissues of C57BL/6J mice, HIIT-trained C57BL/6J mice, HIIT-trained human muscle tissue, and wild mice into aged mice. At the same time, I injected mitochondria from HIIT-trained C57BL/6J and wild mice gastrocnemius tissues into 5-month-old normal C57BL/6J mice. Additionally, mitochondria from each group were also co-cultured with normal HepG2 and D-gal-treated HepG2 cells. ATP and MMP assays were employed to evaluate mitochondrial function. The Wild Mice group exhibited significant differences compared to the C2C12, Young C57 and Trained People groups. Although no significant difference was observed between the Wild Mice and Trained C57 groups, the overall function of the Wild Mice mitochondria was slightly superior (Extend Figures 2A-B). No immune or inflammatory responses were detected 24 hours post-transplant, as indicated by measurements of IL-6, IL-10, and TNF-α (Figure S4A). In the aging model, exercise performance improved across all groups three days after mitochondrial transplantation. The Wild Mice and Trained C57 groups exhibited superior endurance, particularly the Wild Mice group, which demonstrated the best long-term exercise endurance (Extend Figure 2C). In the normal model, only the Wild Mice group showed an increase in exercise capacity (Extend Figure 2I). One week post-transplantation, body weight significantly increased in the aging model, with the Wild Mice group achieving the most substantial recovery (Extend Figures 2D-E). In contrast, no significant weight differences were observed in the normal model (Extend Figures 2J-K). Despite the beneficial effects observed across all transplantation groups, the Wild Mice group demonstrated a more pronounced reduction in oxidative stress among aging mice. The MDA levels in the Wild Mice group were significantly lower compared to those in the Young C57 and Trained People groups (Extend Figures 2F). NOS levels were also lower in the Wild Mice group than in the C2C12, Young C57, and Trained People groups. Although the Trained C57 group exhibited differences from the Young C57 group, it remained inferior to the Wild Mice group (Extend Figure 2G). SOD activity in the Wild Mice group was higher than in the C2C12, Young C57, and Trained C57 groups (Extend Figure 2H), and Wild Mice mitochondria significantly enhanced SOD activity in normal mice (Extend Figure 2M). Furthermore, all mitochondrial transplantation groups improved bioenergetics in normal HepG2 cells, with the Wild Mice and Trained C57 groups exhibiting the most significant increases, particularly the Wild Mice group (Extend Figure 2O). Mitochondrial transplantation restored membrane potential and reduced ROS generation in D-gal-treated HepG2 cells, with the Wild Mice and Trained C57 groups demonstrating a superior ability to alleviate oxidative stress and restore mitochondrial membrane potential (Extend Figures 3A-B).

**Extend Figure3 :**
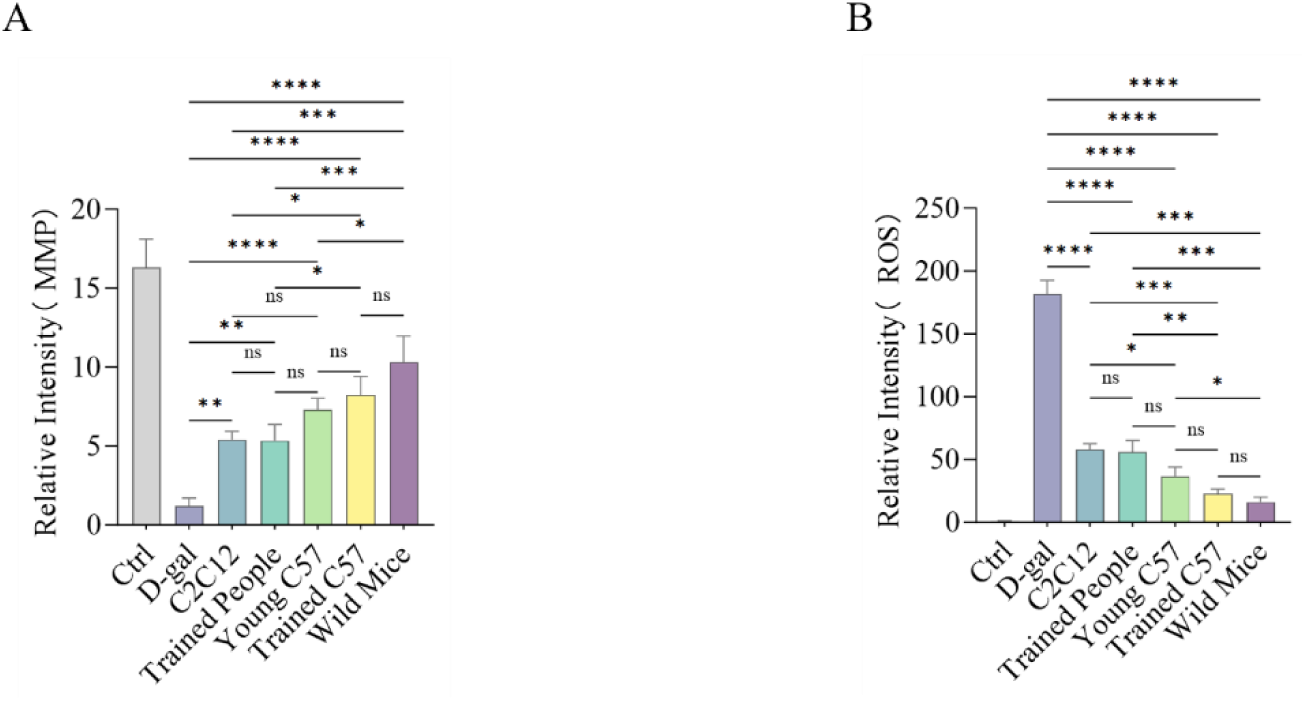
Mitochondrial Transplantation Enhances the Biological Function of HepG2 Cells. (A)The JC-1 statistical analysis method was employed to assess the recovery of mitochondrial membrane potential fluorescence intensity in HepG2 cells treated with D-gal after five different species of mitochondrial transplantation. Untreated cells served as controls, n = 3. (B) Statistical analysis was conducted to evaluate the effects of mitochondrial transplantation from five different species on the fluorescence intensity of reactive oxygen species (ROS) in D-gal-treated HepG2 cells, with untreated cells again serving as controls, n = 3. ns = no statistical significance, * = p<0.05, ** = p<0.01, *** = p<0.001, **** = p<0.0001

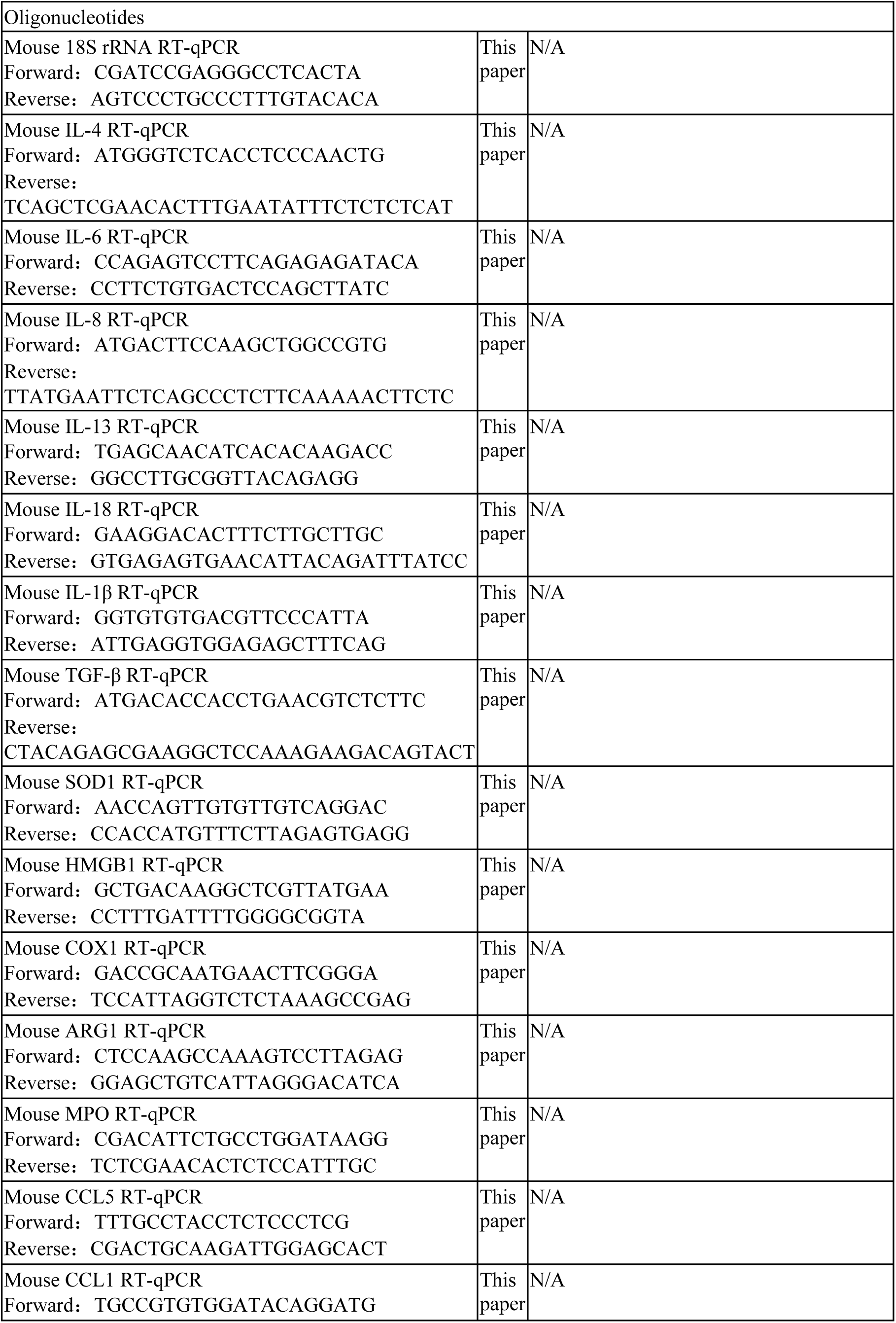
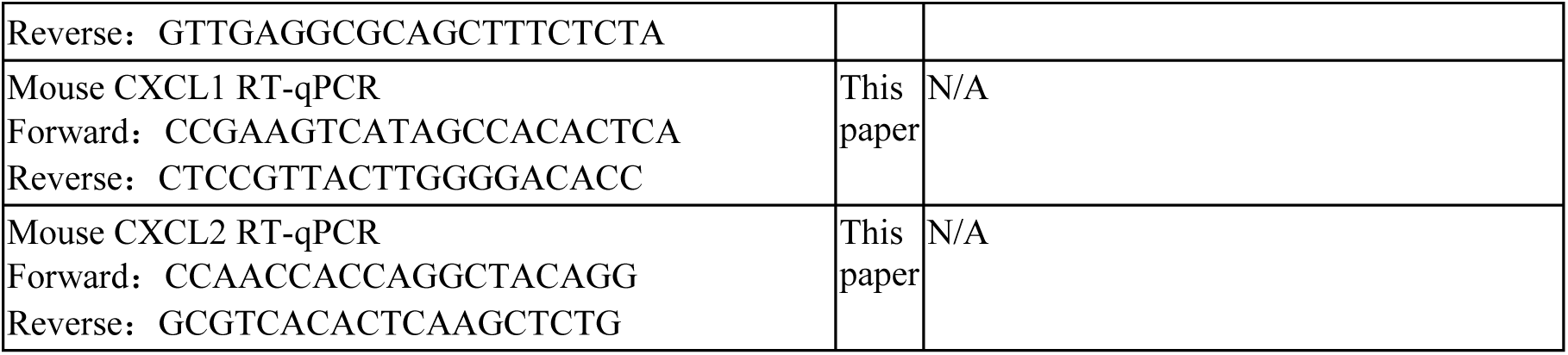

